# RubisCO-encoding methanotrophs elicit a coordinated transcriptional response to carbon dioxide limitation

**DOI:** 10.64898/2026.11.03.764036

**Authors:** Spencer A. Lee, Jessica M. Henard, Yao-Chuan Yu, Calvin A. Henard

## Abstract

The methanotroph *Methylococcus capsulatus* encodes the carbon dioxide (CO_2_)-fixing enzyme ribulose-1,5-bisphosphate carboxylase/oxygenase (RubisCO), which, along with CO_2_ and methane (CH_4_), are required for growth of this bacterium. This unique, dual one-carbon metabolism is present in phylogenetically diverse methanotrophs and is likely widespread in nature, but the connection between organic and inorganic carbon metabolism in these methanotrophs is unclear. Here we used a systems approach to investigate the requisite role of CO_2_ in *M. capsulatus* metabolism and physiology. We show that atmospheric CO_2_ levels are sufficient for *M. capsulatus* cultivation, and that CO_2_ assimilation is linked to cellular redox balance. In the absence of CO_2_, *M. capsulatus* elicits a global transcriptional response that includes induction of RubisCO- and other inorganic carbon assimilation-encoding genes with a concomitant decrease in ribosome biogenesis and nitrogen assimilation genes typical of a bacterial stringent response. This CO_2_ starvation response balances biosynthesis with carbon availability. Genetic and biochemical evidence show that a LysR-type transcriptional regulator (CbbR) activates RubisCO and other CO_2_ assimilation gene transcription via direct promoter binding in response to elevated intracellular RubisCO substrate, ribulose-1,5-bisphosphate, in CO_2_-starved cells. These results offer insights into the essentiality of CO_2_ for *M. capsulatus* metabolism and establish a function for CbbR-family regulators in methanotrophs beyond their canonical association with CO_2_ fixation in autotrophic bacteria.

**Significance:** Methanotrophic bacteria are the primary biological sink of atmospheric methane and represent candidates to develop methane mitigation biotechnologies. Several methanotrophs encode the carbon dioxide-fixation enzyme, RubisCO, enabling them to use carbon dioxide and methane, but the role of carbon dioxide in their metabolism and physiology is not understood. Here, we identified how a RubisCO-encoding methanotroph, *Methylococcus capsulatus*, responds to carbon dioxide starvation. The methanotrophs alter their gene expression to produce proteins that facilitate carbon dioxide capture, a phenotype like that observed in carbon dioxide-starved autotrophic and phototrophic bacteria. These findings show how methanotrophs adapt to changing environmental conditions and emphasize evolutionary relationships between methanotrophs and autotrophs. This knowledge can also guide the use of methanotrophs for reducing climate-active atmospheric gases.

## Introduction

Aerobic methane (CH_4_)-oxidizing bacteria (methanotrophs) play an important role in biogeochemical carbon cycling as one of the primary sinks of CH_4_ in the environment (1–3). Typically low-abundant members of terrestrial soil and freshwater and marine sediment microbial communities, methanotrophs can be enriched in niches with elevated geologically or biologically derived CH_4_ (e.g. CH_4_ seeps, landfills, wetlands) (4–8). In addition to free-living, these bacteria were also recently discovered to be primary components of tree microbiomes and can be found in the mammalian gut (9–11). In these diverse environments, methanotrophs serve as biofilters, capturing CH_4_ before escaping to, or directly from, their atmosphere. The unique capacity to utilize CH_4_ as a carbon and energy source also affords opportunities to leverage methanotrophs for atmospheric CH_4_ abatement and/or biomanufacturing valuable products using CH_4_ as an inexpensive feedstock (12, 13).

The ability of methanotrophs to utilize CH_4_ as a carbon and energy source is engendered by enzymes that oxidize CH_4_ through a multi-step pathway to carbon dioxide (CO_2_), thereby generating reducing equivalents for ATP production via oxidative phosphorylation (14). A fraction of partially oxidized CH_4_ intermediates are assimilated into biomass via two primary C1 metabolic pathways that occur in phylogenetically distinct methanotrophs: formaldehyde via the ribulose monophosphate (RuMP) cycle in Gammaproteobacteria methanotrophs or formate via the serine cycle in Alphaproteobacteria methanotrophs (15). Alternatively, the Verrucomicrobiota methanotrophs completely oxidize all CH_4_ to CO_2_ and re-assimilate the inorganic carbon into biomass through the Calvin-Benson-Basham (CBB) cycle (16).

Despite divergent C1 assimilation metabolic pathways, all methanotrophs must oxidize some portion of CH_4_ completely to CO_2_ to generate sufficient reducing equivalents for growth or maintenance (15); thus, all methanotrophs evolve CO_2_. However, many methanotrophs can reassimilate CO_2_ using carboxylases. Alphaproteobacterial methanotrophs assimilate equimolar CO_2_ per CH_4_ oxidized in the phosphoenolpyruvate carboxylase step of the serine cycle (17).

Carbon tracing experiments show that up to 70% of the Alphaproteobacteria methanotroph, *Methylosinus trichosporium* OB3b, biomass is derived from exogenous CO_2_ (18). Further, a subset of Gammaproteobacteria and Alphaproteobacteria methanotrophs encode RubisCO (19–21), presumably to capture CH_4_-derived CO_2_ and enhance metabolic flexibility.

*Methylococcus capsulatus* was one of the first methanotrophs to be isolated and has served as a model CH_4_-oxidizing bacterium to shape the current understanding of methanotrophy (22–24). *M. capsulatus* is a Gammaproteobacteria methanotroph that utilizes the RuMP cycle for CH_4_ assimilation (25) and also encodes RubisCO (26, 27), similar to other closely related Type Ib methanotrophs in the genera *Methylocaldum*, *Methyloterricola*, *Methylomagnum*, and *Methylogaea* (2, 28–30). These methanotrophs also encode phosphoribulokinase and the other CBB cycle enzymes that have overlapping roles in the RuMP cycle (24). Prior CO_2_ isotopic tracing experiments indicate *M. capsulatus* RubisCO is active and that ∼20% of its biomass is derived from CO_2_ (19). Further, CO_2_ and RubisCO are essential for *M. capsulatus* cultivation (19). Notably, attempts to culture *M. capsulatus* autotrophically with hydrogen or formate as energy sources have yielded inconsistent results, with one study demonstrating successful autotrophic growth with hydrogen supplementation only on solid medium (31). Collectively, these prior results underscore a close link between CH_4_ and CO_2_ metabolism in this methanotroph.

Here we used a systems-based approach to investigate the role of CO_2_ in *M. capsulatus*physiology, which revealed significant alterations to both the transcriptome and metabolome in response to limited CO_2_ availability. Several dissolved inorganic carbon metabolic enzymes including RubisCO, carbonic anhydrases, and SulP family transporters were upregulated in response to CO_2_ starvation. We show that transcription of RubisCO and a putative carbonic anhydrase is directly regulated by a LysR-type transcriptional regulator that is allosterically modulated by the RubisCO substrate ribulose-1,5-bisphosphate that quickly accumulates in cells starved for CO_2_. Like cyanobacteria and other autotrophic bacteria, the *M. capsulatus* CO_2_ starvation response enables this bacterium to scavenge limited CO_2_ and maintain its unique methanoautotrophic metabolism. These results provide insight into the metabolic versatility of *M. capsulatus* and other RubisCO-encoding methanotrophs in nature. Additionally, the CO_2_ starvation response and its regulator identified here reveal potential engineering targets to improve carbon capture and conversion for methanotroph-based bioremediation and biotechnology applications.

## Results

### *M. capsulatus* elicits a transcriptional response to CO_2_ limitation

We previously identified CO_2_ as requisite for *M. capsulatus* growth in a bioreactor with continuous gas sparging (19). Here, we evaluated methanotrophic growth on 20% CH_4_ with increasing concentrations of CO_2_ in the supplied gas stream to identify the minimum CO_2_ concentration required for growth. Consistent with prior results (19), *M. capsulatus* displayed optimal growth with a 20% CH_4_/2% CO_2_ in air gas mixture but did not grow when supplied with a 20% CH_4_ in air (0.032% CO_2_) mixture (Fig. 1a), presumably because bacteria-evolved CO_2_ is stripped. *M. capsulatus* was able to grow if the CO_2_ was raised to the atmospheric level (0.04%), displaying a decreased growth rate but reaching a similar terminal OD_600_ compared to the cultures with saturating 2% CO_2_ levels (Fig. 1a). We measured CO_2_ in the effluent gas streams of inoculated cultures compared to uninoculated growth medium supplied with 20% CH_4_ in air (0.032% CO_2_), which showed that the methanotrophs (∼ 1 x 10^9^ CFU) were evolving CO_2_ up to 24 h in the absence of detectable growth (Fig. 1b). Cells also remain viable without CO_2_ for up to 96 h (Fig. 1c), which aligns with the ability of the CO_2_-starved cells to recover growth if CO_2_ is reintroduced to the supplied gas stream (19). The absence of RubisCO-dependent CO_2_ assimilation led to an increase in the NADPH/NADP+ ratio due to an elevated reduced NADPH pool (Fig. 1d), but the total intracellular NADP(H) and ATP concentrations were similar between CO_2_-replete and -deplete cells 6 h following CO_2_ starvation (Fig. 1e and 1f). These data show that *M. capsulatus* requires only atmospheric levels of CO_2_ for growth and that CO_2_-starved cultures are unable to expand but remain metabolically active.

**Figure 1.**
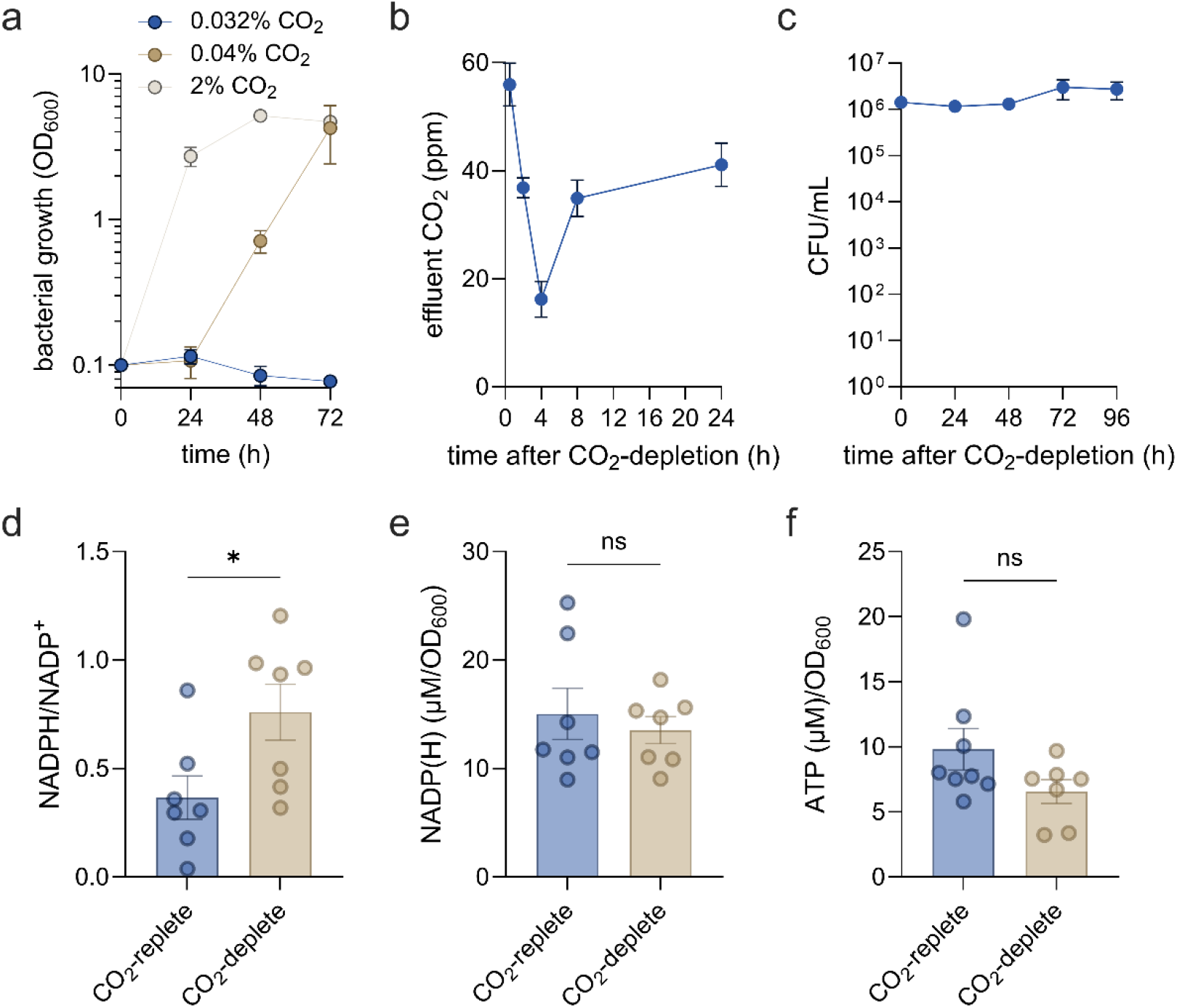
Atmospheric CO_2_ levels support *M. capsulatus* growth in a bioreactor with continuous gas sparging. a) *M. capsulatus* growth in a custom bioreactor with continuous gas sparging of 20% CH_4_ and increasing CO_2_ levels balanced with air. The baseline gas mixture (20% CH_4_/80% air contains 0.032% CO_2_ measured by gas chromatography. b) CO_2_ in the effluent gas stream evolved from *M. capsulatus* cultures (OD_600_ = 0.1) after additional CO_2_ was removed from the gas stream (CO_2_-deplete). Values represent values above baseline (uninoculated medium control). c) *M. capsulatus* viability (colony forming units, CFU) over time after CO_2_ removal from the gas stream. d) The NADPH/NADP^+^ ratio, (e) total NADP(H), and (f) ATP quantified in extracts from *M. capsulatus* cultivated with continuous 20% CH_4_/2% CO_2_ in air (unstarved, blue bars) or 5 h after removal of CO_2_ from the gas stream (CO_2_-deplete, tan bars). The data represent the mean ± SEM (n=4-7) from at least two independent experiments. \**p* ≤ 0.05 from unpaired *t*-test; ns, not statistically significant.

To identify potential *M. capsulatus* physiological responses to CO_2_ starvation, we compared transcription in bioreactor cultures supplied with 20% CH_4_/2% CO_2_ in air (unstarved) to those after CO_2_ was removed from the sparge gas stream (CO_2_-deplete) (Fig. 2a). RNA sequencing identified 388 differentially expressed genes (DEGs) 1 h following CO_2_ depletion with 83 genes upregulated (*p* ≤ 0.001, log_2_FC ≥ 1) and 305 genes downregulated (*p* ≤ 0.001, log_2_FC ≤ -1) (Fig. 2b, Table 1, and Table S1). The global transcriptional response expanded to 568 DEGs (*p* ≤ 0.001, │log_2_FC│ ≥ 1) by 3 h post CO_2_ starvation with 198 genes up-regulated and 370 genes down-regulated (Fig. S1a and Table S2). Many of the downregulated genes encode proteins involved in nitrogen transport and metabolism and protein synthesis (Fig. S1b and Table S3), consistent with the guanine tetraphosphate (ppGpp) alarmone-dependent stringent response that is triggered by carbon or nitrogen starvation in phylogenetically diverse bacteria (32, 33). Additionally, transcription of several core CH_4_ assimilation/dissimilation pathway genes decreased under CO_2_-deplete conditions, including particulate methane monooxygenase (MCA2854); pQQ-dependent methanol dehydrogenase (MCA0079) formaldehyde activating enzyme (MCA2866), and several enzymes from the tetrahydromethanopterin CH_4_ assimilation pathway (MCA3018-MCA3019), multiple formate dehydrogenase isozymes (MCA1210, MCA1391), enzymes from the ribulose monophosphate pathway (MCA0024-MCA0025, MCA1251, MCA1950) and key enzymes in the Entner-Doudoroff pathway (MCA0037-MCA0038) (Fig. S1c, Table S4). Conversely, genes involved in inorganic carbon metabolism, including the operon encoding RubisCO and the RubisCO activase (MCA2743-2746), putative HCO_3_^-^ transporters (MCA2472 and MCA1443) and a carbonic anhydrase (MCA1442), were significantly upregulated in response to CO_2_ starvation (Fig. S1c and Table S4). Notably, a gene of unknown function (MCA2457) was the most induced (log_2_FC = 4.73), and, together with the RubisCO operon genes, comprised the most abundant mRNA transcripts in the transcriptome (Fig. 2b). The MCA2457 gene encodes a 156 amino acid protein with a nuclear transport factor 2 (NTF2)-like fold found in SgcJ/EcaC family oxidoreductases and the recently identified iota subclass of carbonic anhydrases (ι-CA) (34–36). Interestingly, BLASTp analysis of MCA2457 showed it has significant amino acid identity to proteins annotated as SgcJ/EcaC family oxidoreductases in the genera of the Type Ib methanotrophs *Methylococcus Methylocaldum*, and *Methylospira* (those that typically encode RubisCO) and *Mesorhizobium* spp. Further, MCA2457 is predicted by SignalP6.0 to contain a 24 amino acid N-terminal Sec secretion signal sequence, so the protein is likely secreted into the methanotroph periplasm as a 15.4 kilodalton mature protein after signal peptide cleavage. These results support that MCA2457 may encode a periplasmic ι-CA homolog that could facilitate inorganic carbon capture.

**Figure 2.**
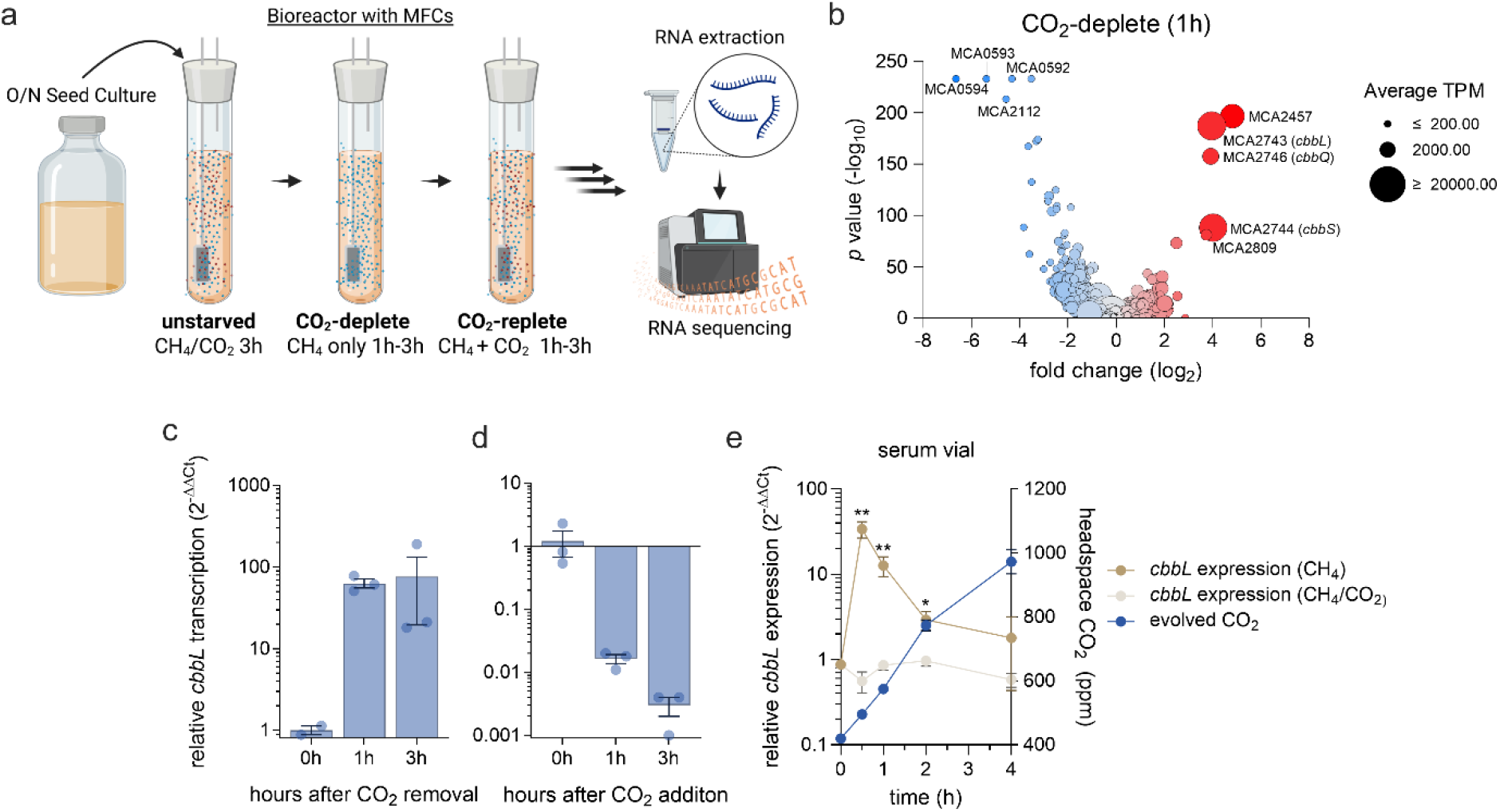
*M. capsulatus* elicits a global transcriptional response to CO_2_ starvation. a) Schematic of the bioreactor experimental set-up wherein *M. capsulatus* is sub-cultured from an overnight serum vial seed culture into a bioreactor Kimax bubble column and cultivated with continuous gas sparging of 20% CH_4_/2% CO_2_ in air for 3 h followed by removal of the CO_2_ from the gas stream (20% CH_4_ in air, CO_2_-deplete). Alternatively, cells were sub-cultured from serum vials and incubated under CO_2_-deplete conditions for 3 h followed by the addition of CO_2_ to the gas stream (CO_2_-replete). b) Volcano plot of differential expressed genes 1h after CO_2_-depletion compared to unstarved controls. Red corresponds to upregulated genes and blue to downregulated genes with color intensity correlated to DEG log_2_ fold change. Dot size correlates to mean gene transcripts per million (TPM). c-d) RT-qPCR analysis of the gene (MCA2743, *cbbL*) encoding the RubisCO large subunit under (c) CO_2_-deplete and (d) CO_2_-replete cultivation conditions compared to unstarved or CO_2_-deplete controls (0 h), respectively. e) RT-qPCR analysis of *cbbL* expression following *M. capsulatus* inoculation into sealed serum vials with atmospheres containing 20% CH_4_ in air with (beige) or without (dark tan) the addition of 8% CO_2_ added to the headspace. *cbbL* expression is negatively correlated to *M. capsulatus* CH_4_-derived CO_2_ evolution into the vial headspace (blue). Data in b are based on RNA sequencing from 3 biological replicates for each condition, and data in c-e represent the mean ± SEM (n=3). \**p* ≤ 0.05, \*\**p* ≤ 0.01 from unpaired *t*-tests.

**Table 1.** The most differentially expressed genes (*p* ≤ 0.001, log_2_ ≥ 1.5 or log_2_ ≤ -3) in *M. capsulatus* 1 h after CO_2_ removal from the bioreactor sparge gas (CO_2_-deplete).

| **Name** | **Differential Expression Log2 Ratio** | **Differential Expression *p*-value** | **locus_tag** | **old_locus_tag** |
|---|---|---|---|---|
| **nuclear transport factor 2 family protein CDS** | 4.33 | 1.73E-197 | MCA_RS12055 | MCA2457 |
| **form I ribulose bisphosphate carboxylase large subunit CDS** | 3.76 | 4.18E-188 | MCA_RS13440 | MCA2743 |
| **ribulose bisphosphate carboxylase small subunit CDS** | 3.60 | 7.15E-89 | MCA_RS13445 | MCA2744 |
| **CbbQ/NirQ/NorQ/GpvN family protein CDS** | 3.51 | 3.19E-158 | MCA_RS13450 | MCA2746 |
| **hypothetical protein CDS** | 3.40 | 7.22E-82 | MCA_RS13755 | MCA2809 |
| **hypothetical protein CDS** | 2.66 | 2.00E-22 | MCA_RS01695 |  |
| **cellulose binding domain-containing protein CDS** | 2.05 | 4.65E-74 | MCA_RS13455 | MCA2747 |
| **SulP family inorganic anion transporter CDS** | 1.86 | 2.98E-41 | MCA_RS07105 | MCA1443 |
| ***dnaK* CDS** | 1.80 | 8.11E-15 | MCA_RS09120 | MCA1856 |
| **ammonium transporter CDS** | 1.79 | 1.19E-33 | MCA_RS02425 | MCA0490 |
| **carbonic anhydrase CDS** | 1.60 | 8.28E-11 | MCA_RS07100 | MCA1442 |
| **hypothetical protein CDS** | 1.59 | 3.86E-15 | MCA_RS16225 |  |
| **HNH endonuclease CDS** | 1.57 | 7.19E-30 | MCA_RS12710 | MCA2596 |
| **lasso peptide biosynthesis B2 protein CDS** | 1.55 | 6.58E-15 | MCA_RS15615 | MCA0888 |
| **GFA family protein CDS** | 1.55 | 3.43E-12 | MCA_RS02450 | MCA0499 |
| **hypothetical protein CDS** | 1.51 | 7.48E-29 | MCA_RS01400 | MCA0286 |
| **Rieske 2Fe-2S domain-containing protein CDS** | -3.06 | 6.68E-41 | MCA_RS02905 | MCA0586 |
| ***nifA* CDS** | -3.41 | 3.79E-175 | MCA_RS03775 | MCA0764 |
| ***nifL* CDS** | -3.54 | 1.80E-133 | MCA_RS03770 | MCA0763 |
| **nitrate reductase CDS** | -3.55 | 4.96E-173 | MCA_RS02930 | MCA0590 |
| ***nirD* CDS** | -3.73 | 4.65E-63 | MCA_RS02935 | MCA0591 |
| ***glnK* CDS** | -3.85 | 3.26E-89 | MCA_RS10450 | MCA2137 |
| **ammonium transporter CDS** | -3.92 | 4.24E-234 | MCA_RS10445 | MCA2136 |
| **alginate export family protein CDS** | -4.00 | 4.84E-168 | MCA_RS02910 | MCA0587 |
| ***nirB* CDS** | -4.51 | 0 | MCA_RS02940 | MCA0592 |
| **porin CDS** | -5.00 | 6.29E-214 | MCA_RS10325 | MCA2112 |
| **bifunctional protein-serine/threonine kinase/phosphatase CDS** | -5.69 | 0 | MCA_RS02945 | MCA0593 |
| **MFS transporter CDS** | -7.07 | 0 | MCA_RS02950 | MCA0594 |

The CO_2_ starvation response was reversed within 1 h upon reintroduction of 2% CO_2_ to the CO_2_-deplete cultures (Fig. S1d, Table S5), with previously starved cells having a similar transcriptome to unstarved controls by 3 h after CO_2_ addition (Fig. S1e-f, Tables S6 and S7). RT-qPCR analysis showed that the *cbbL* gene (MCA2743) encoding the RubisCO large subunit was induced 100-fold following CO_2_ starvation and returned to baseline levels after CO_2_ was introduced to starved cultures (Fig. 2c-d), validating the RNA-seq data. *cbbL* transcription was also induced following *M. capsulatus* inoculation into serum vials with a 20% CH_4_ in air atmosphere but was quenched as CH_4_ oxidation-derived CO_2_ accumulates in the headspace (> 500 ppm CO_2_) (Fig. 2e), indicating the CO_2_ starvation response is also induced upon inoculation under these standard methanotroph growth conditions. This phenotype was abolished if exogenous CO_2_ was added to the serum vial (20% CH_4_/8% CO_2_ in air) at inoculation (Fig. 2e), an atmospheric mixture that decreases the *M. capsulatus* lag phase in serum vials (19). To determine whether this response is generalizable in RubisCO-encoding methanotrophs, we also evaluated *cbbL* transcription in another Type Ib methanotroph, *Methylocaldum gracile* (29), which was significantly induced in response to CO_2_ starvation (Fig. S2).

Given the significant decrease in *M. capsulatus* translational machinery transcription in response to CO_2_ starvation, we questioned whether the observed increase in RubisCO transcription would positively correlate to an increase in RubisCO protein. To address this question, we developed an *M. capsulatus* strain expressing RubisCO with a 3xFlag epitope tag on the N-terminus of the large CbbL subunit. This strain was subjected to CO_2_-depletion and cells were collected over time for assessing RubisCO expression via Western blotting, which showed a 2.6-fold increase in 3xFlag-CbbL 24 h following CO_2_ starvation (Fig. 3a). To evaluate whether the increase in RubisCO mediates enhanced CO_2_ uptake after CO_2_ starvation, we quantified the rate of ^13^CO_2_ assimilation by previously CO_2_-starved cells using LC-MS/MS isotopic metabolomics. ^13^C incorporation into the RubisCO product 3-phosphoglycerate (3PG) and regeneration of the ribulose-1,5-bisphosphate (RuBP) substrate in CO_2_-deplete cells was 2-fold higher compared to unstarved cells (Fig. 3b), indicating not only higher rates of CO_2_ assimilation consistent with the elevated RubisCO levels (Fig. 3a), but also increased carbon flux through the *M. capsulatus* CBB cycle.

**Figure 3.**
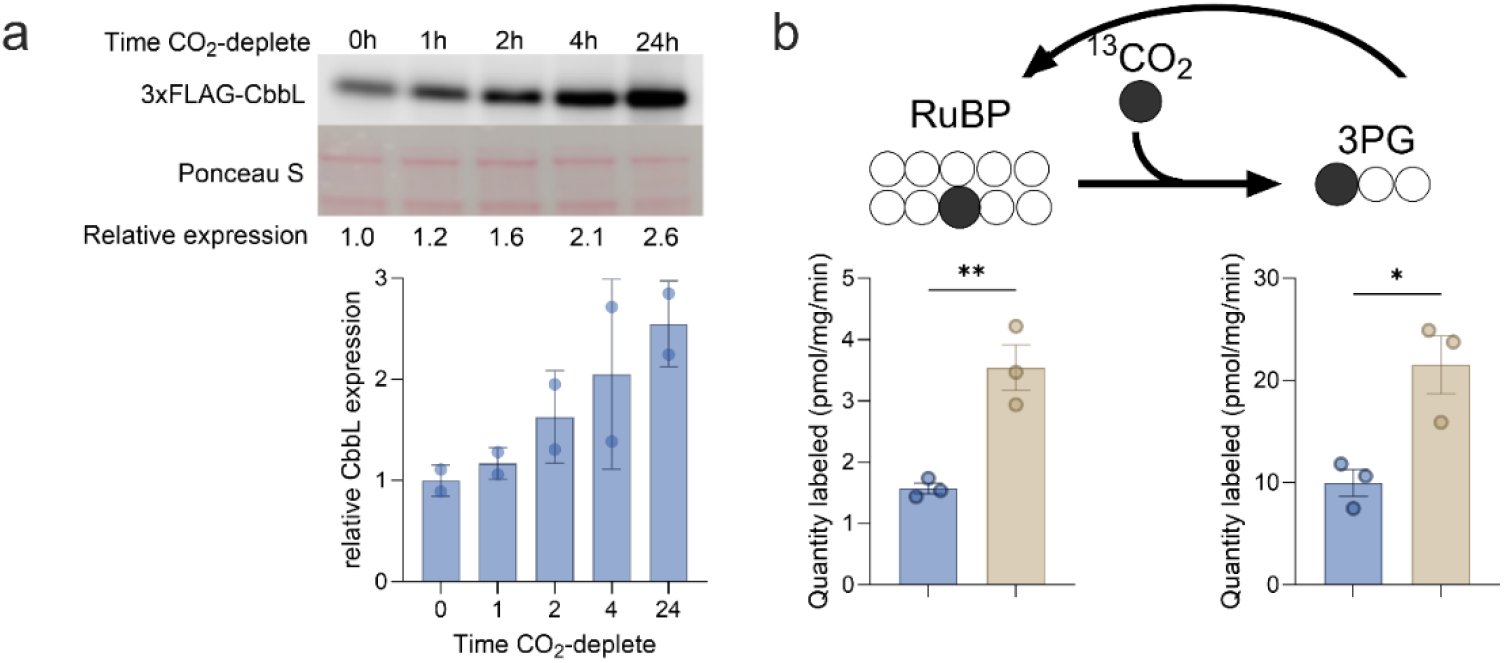
The CO_2_ starvation response enhances *M. capsulatus* inorganic carbon assimilation and conversion upon reintroduction. a) Immunoblot of 3xFlag-CbbL in *M. capsulatus* cell lysates over time following CO_2_ starvation. Ponceau S staining of the membrane was used as a loading control. Densitometric quantification of 3xFlag-CbbL normalized to Ponceau S. Values are expressed as fold change relative to unstarved controls (mean ± SD) from two independent experiments. b) Quantification of intracellular ribulose-1,5-bisphosphate (RuBP) and 3-phosphoglycerate (3PG) ^13^C-labeled isotopologues after unstarved (blue bars) or CO_2_-starved (4h, tan bars) cells were incubated with ^13^CO_2_ for 5 minutes. Data represent the mean ± SEM (n=3). \**p* ≤ 0.05, \*\**p* ≤ 0.01 from unpaired *t*-tests.

### A targeted CRISPRi screen identifies the CO_2_ starvation response genes important for *M. capsulatus* fitness during CH_4_ and CO_2_-dependent growth

We next leveraged recently developed methanotroph functional genomic CRISPRi tools and workflows to assess the potential role of the genes differentially expressed in response to CO_2_ starvation (37). A library consisting of 6,220 sgRNAs, 15 sgRNAs per the 388 differentially expressed genes (Table S1, *p* ≤ 0.001, │log_2_FC│ ≥ 1) and 400 non-targeting control sgRNAs was designed and synthesized via massively parallel oligo synthesis (Table S8). The pooled sgRNA library was cloned downstream of the anhydrotetracycline-inducible P*_tet_* promoter in the broad-host-range plasmid pCAH01 and then transferred via conjugation to an *M. capsulatus* strain with a chromosomally integrated P*_tet_*-dCas9 cassette to enable inducible CRISPRi-dependent transcription repression. This methanotroph targeted sgRNA library was cultivated in serum vials with an atmosphere of 20% CH_4_ in air or 20% CH_4_/8% CO_2_ with and without CRISPRi induction to identify the CO_2_ starvation-responsive genes essential for *M. capsulatus* growth. As expected based on prior observations (19), CO_2_ supplementation significantly decreased the lag phase/boosted growth of the targeted library in both induced and uninduced samples (Fig. 4a). The sgRNA library composition in CRISPRi induced and noninduced samples 72 h after selection was quantified by Oxford Nanopore Technologies sgRNA amplicon sequencing and mapping to the synthesized sgRNA library. We identified 98.7% (6,138/6,220) of the synthesized sgRNAs in the sample sequences, with every target gene having multiple corresponding sgRNAs represented (Tables S9 and S10). Comparing the sgRNA library composition between CRISPRi induced vs uninduced samples from the library cultivated without CO_2_ supplementation, conditions when the CO_2_ starvation response is induced in early lag phase (Fig. 2e), showed that 583 sgRNAs (9.5%) were depleted (*p* ≤ 0.001, log_2_ fitness score ≤ 1) and 23 sgRNAs (0.4%) were enriched (*p* ≤ 0.001, log_2_ fitness score ≤ 1) after 72 h of selection (Fig. 4b outer ring, Table S9), which correspond to a fitness defect or advantage phenotype associated with transcription repression of the sgRNA target gene.

**Figure 4.**
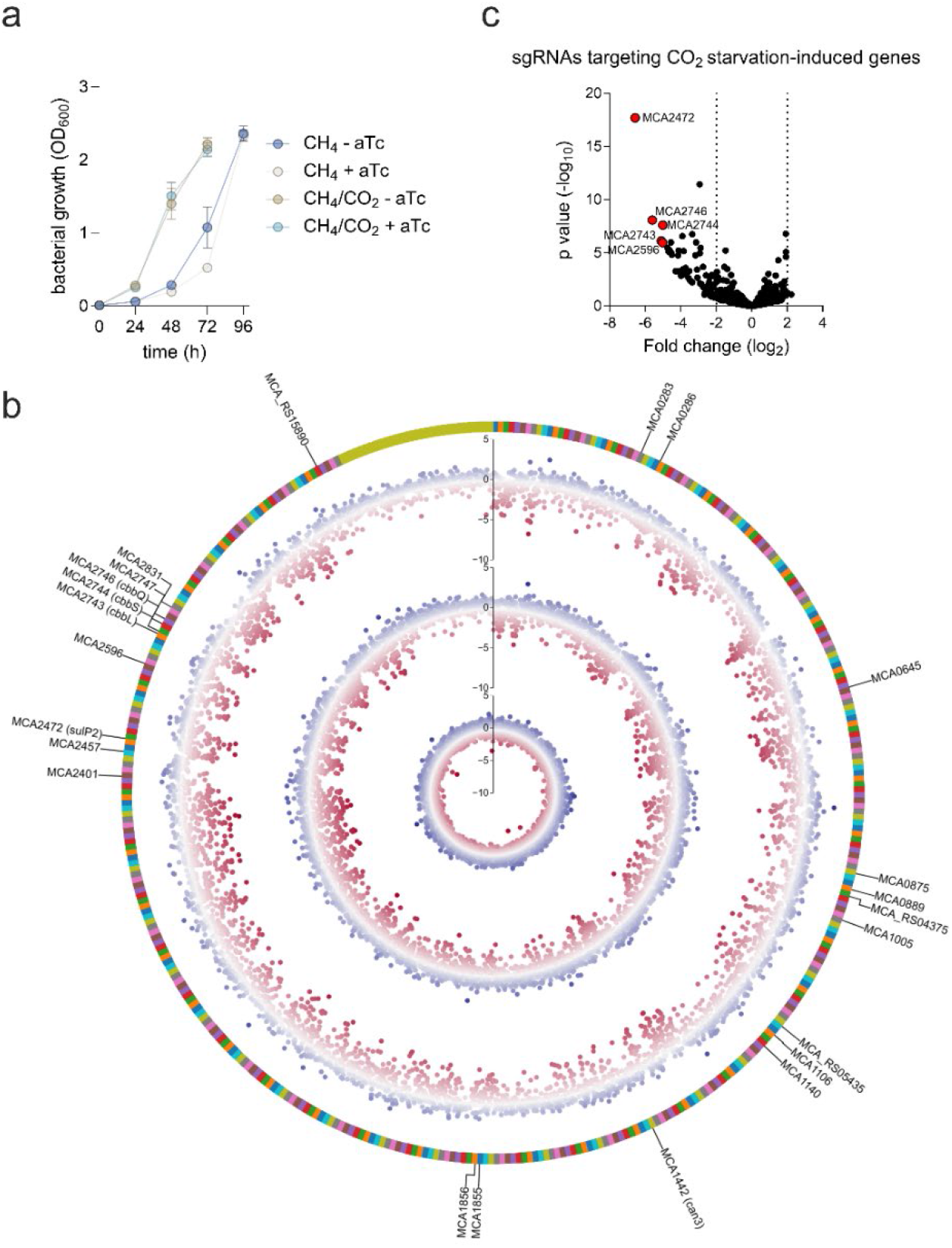
A targeted CRISPRi library screen identifies key players of the *M. capsulatus* CO_2_ starvation response. a) Cultivation of an *M. capsulatus* CRISPRi library targeting the 388 genes differentially expressed in response to CO_2_ starvation in serum vials with 20% CH_4_ in air or 20% CH_4_/8% CO_2_ in air with (+ aTc) and without (-aTc) sgRNA and dCas9 induction to repress target gene transcription. b) Circos plot depicting the 6,220 sgRNA library differential fitness scores (dots) calculated by comparing sgRNA abundances for induced vs uninduced after 72 h of growth with CH_4_ (second ring), or CH_4_ supplemented with CO_2_ (third ring). Differential sgRNA fitness scores comparing CH_4_ to CH_4_/CO_2_ with CRISPRi induction (inner ring). The outer ring represents the 388 genes targeted in the screen (multicolor) and the non-targeting sgRNAs (mustard). In the differential fitness score rings, sgRNAs with negative fitness scores are red and those with positive fitness scores are blue with color intensity correlated to the log_2_ fold change. c) Differential fitness of sgRNAs from serum vial cultures without CO_2_ supplementation targeting those genes induced (log_2_FC ≥ 1 and *p* ≤ .001) by CO_2_-starvation identified in Fig. 2. Data in (a) represent the mean ± SEM (n=4). Data in b and c are based on sgRNA amplicon sequencing from 4 biological replicates for each condition

Several sgRNAs targeting the 83 genes induced (*p* < 0.001, log_2_FC > 1) in response to CO_2_ starvation were significantly depleted during growth with CH_4_ without CO_2_ supplementation (Table 2 and Fig. 4c), underscoring the important role of the CO_2_ starvation response genes for *M. capsulatus* fitness and growth. Interestingly, one of the most depleted sgRNAs targets MCA2472 (log_2_FC = -6.57), which is annotated to encode a SulP bicarbonate transporter that may be needed for *M. capsulatus* inorganic carbon uptake and utilization. sgRNAs targeting the RubisCO and RubisCO activase operon genes were also significantly depleted (Fig. 4c; MCA2546 (*cbbQ*), log_2_FC = -5.60; MCA2743 (*cbbL*), log_2_FC = -5.11; MCA2744 (*cbbS*), log_2_FC = -5.01). An sgRNA targeting a β-CA-encoding gene (MCA1442) was also one of the most depleted (log_2_FC = -4.55). This CA was previously shown to be essential for *M. capsulatus* growth and, when overexpressed, can improve carbon conversion efficiency in this methanotroph (38). Additionally, repressing transcription of the putative CA encoded by MCA2457 resulted in a fitness defect (log_2_FC = -3.91).

**Table 2.** Genes induced in response to CO_2_ starvation with the most differential sgRNA fitness scores (*p* ≤ 0.001, log_2_ ≤ -3) identified in a targeted CRISPRi library screen cultivated without CO_2_ supplementation.

| **Gene Description** | **sgRNA Target Gene** | **sgRNA Differential Fitness Log2 Ratio** | **sgRNA Differential Fitness *p*-value** |
|---|---|---|---|
| **SulP family inorganic anion transporter CDS** | MCA2472 | -6.57 | 2.00E-18 |
| **CbbQ RubisCO activase** | MCA2746* | -5.60 | 8.55E-9 |
| **RubisCO large subunit** | MCA2743* | -5.11 | 7.51E-07 |
| **HNH endonuclease** | MCA2596 | -5.03 | 1.09E-6 |
| **RubisCO small subunit** | MCA2744* | -5.01 | 7.51E-7 |
| **hypothetical protein** | MCA0286* | -4.75 | 3.39E-6 |
| **carbonic anhydrase** | MCA1442 | -4.55 | 9.34E-6 |
| **DnaK chaperone** | MCA1856 | -4.50 | 1.54E-5 |
| **hypothetical protein** | MCA0645 | -4.16 | 1.75E-5 |
| **formate hydrogenlyase** | MCA1140 | -4.13 | 6.29E-6 |
| **inorganic phosphate transporter** | MCA1005* | -4.12 | 3.47E-4 |
| **nuclear transport factor 2 family protein** | MCA2457 | -3.91 | 2.73E-07 |

\*additional sgRNAs targeting the same gene with *p* ≤ 0.001, log<sub>2</sub> ≤  -3 not shown.

The amount of sgRNAs with significant negative fitness scores (*p* ≤ 0.001, log_2_FC ≤ 1) decreased in cultures supplemented with CO_2_ (530 sgRNAs, 8.6%; Fig 4b middle ring, Table S10), suggesting that transcription repression of some genes has a more profound negative effect on fitness in the absence of CO_2_ supplementation. Indeed, transcription repression of 14 genes resulted in a more severe fitness defect without CO_2_ supplementation (sgRNA fitness scores *p* ≤ 0.001, log_2_FC ≤ 2 comparing CH_4_-to CH_4_/CO_2_-growth; Fig. 4b, inner ring, Table 3, Table S11). Notably, in both datasets, many, but not all, of the sgRNAs targeted the same gene had similar differential fitness scores while 95% of the non-targeting control sgRNAs were similar between CRISPRi induced vs uninduced conditions, supporting the strength of the CRISPRi screening in identifying target genes. Taken together, the decreased fitness associated with repressing genes encoding a bicarbonate transporter, CAs, and RubisCO suggest that a putative carbon concentrating mechanism like that in autotrophic bacteria may be active and required for efficient *M. capsulatus* cultivation.

**Table 3.** The *M. capsulatus* genes with the most differential sgRNA fitness scores (*p* ≤ 0.001, log_2_ ≤ -2 or log_2_ ≥ 2) between methanotroph targeted CRISPRi library serum vial cultures supplied with 20% CH_4_ in air or with 20%CH_4_ and 8% CO_2_ in air.

| Name | sgRNA Target Gene | sgRNA Differential Fitness Log2 Ratio | sgRNA Differential Fitness *p*-value |
|---|---|---|---|
| **CbbQ/NirQ/NorQ/GpvN family protein** | MCA2746* | -3.79 | 1.03E-10 |
| **MoxR family ATPase** | MCA1525 | -3.67 | 2.98E-10 |
| **phosphoserine phosphatase SerB** | MCA1267 | -3.20 | 1.12E-11 |
| **UMP kinase** | MCA0569* | -2.58 | 0.00000706 |
| **30S ribosomal protein S2** | MCA0567 | -2.47 | 0.000178279 |
| **50S ribosomal protein L4** | MCA2371 | -2.39 | 0.000237946 |
| **hypothetical protein** | MCA0193 | -2.35 | 0.000000186 |
| **homocitrate synthase** | MCA0254 | -2.31 | 0.00000577 |
| **2,3-bisphosphoglycerate-independent phosphoglycerate mutase** | MCA0753 | -2.30 | 0.000237736 |
| **cytochrome c** | MCA2401 | -2.20 | 0.0000112 |
| **50S ribosomal protein L5** | MCA2360 | -2.15 | 0.0000815 |
| **molecular chaperone DnaK** | MCA1856 | -2.08 | 3.17E-08 |
| **TolC family protein** | MCA2262 | -2.06 | 0.00096727 |
| **TonB-dependent receptor** | MCA2321 | -2.02 | 0.000138686 |
| **Fe-S cluster assembly scaffold IscU** | MCA0248 | 2.01 | 8.92E-10 |
| **methanol/ethanol family PQQ-dependent dehydrogenase** | MCA0779 | 2.04 | 7.43E-10 |
| **hypothetical protein** | MCA0286 | 2.17 | 0.000112611 |
| **nitrate reductase** | MCA0590 | 2.20 | 8.55E-06 |
| ***nif*-specific transcriptional activator NifA** | MCA0764 | 2.24 | 0.000472517 |
| **S8 family serine peptidase** | MCA0875 | 2.32 | 2.80E-05 |
| **endonuclease III** | MCA2899 | 2.48 | 3.28E-07 |
| **histidine kinase** | MCA0778 | 2.55 | 2.33E-07 |
| **30S ribosomal protein S4** | MCA2348 | 2.70 | 7.40E-06 |

\*duplicate sgRNAs with *p* ≤ 0.001, log<sub>2</sub> ≤  -2 not shown.

### A LysR-type transcriptional regulator mediates the CO_2_ starvation response in *M. capsulatus*

The LysR type transcriptional regulator (LTTR) CbbR (CBB cycle transcriptional regulator) orchestrates responses to CO_2_ availability in an array of chemo- and photo-autotrophic bacteria (39–43). We identified a CbbR homolog in the *M. capsulatus* genome, encoded by MCA3039 with 43.47% amino acid identity to the *Synechococcus elongatus* PCC7942 CCM regulating protein CmpR (PDB:5Z49) as a candidate regulator of the CO_2_ starvation response (Fig. S3a). Interestingly, in contrast to the typical genomic context of characterized CbbR genes that are colocalized with RubisCO genes, MCA3039 is divergently positioned next to a core operon encoding key enzymes of the RuMP cycle, including transketolase, fructose bisphosphate aldolase, hexulose-1-phosphate synthase and isomerase, and transaldolase (Fig. S3b).

An *M. capsulatus* CbbR-deficient knock-out strain (Δ*cbbR*::Kn^R^) exhibited a growth defect compared to a wild-type control in a bioreactor with a 20% CH_4_/0.04% CO_2_ in air gas stream (Fig. 5a), and in serum vials supplied with 20% CH_4_ in air (Figure 5b). In contrast, Δ*cbbR*::Kn^R^ displayed comparable growth to wild-type with continuous 20% CH_4_/2% CO_2_ in air supply (Fig. 5a). This high CO_2_-requisite phenotype suggests that CbbR may play a role in transcriptional regulation of *M. capsulatus* CO_2_ metabolism. Indeed, RT-qPCR analysis showed that induction of both the MCA2457 gene and the RubisCO large subunit *cbbL* gene (MCA2743) in response to CO_2_ starvation was abolished in the Δ*cbbR*::Kn^R^ strain (Fig. 5c-d), supporting that the MCA3039 LTTR is a transcriptional activator of RubisCO and the putative CA encoded by MCA2457. We determined the CbbR regulon by comparing transcription responses of the Δ*cbbR*::Kn^R^ strain during CO_2_-replete and-deplete growth (Fig. 5f and Table S12 and S13). Comparing the DEGs in response to CO_2_ starvation between the Δ*cbbR*::Kn^R^ strain and wild-type indicated that the CbbR regulon is discrete, consisting of only the RubisCO and MCA2457 genes (Fig.5g and Table S14). Notably, 115 genes were induced in response to CO_2_ starvation in Δ*cbbR*::Kn^R^ (Fig. 5f-g, Tables S12 and S13), suggesting other regulators, together with CbbR, likely coordinate the global CO_2_ starvation response.

**Figure 5.**
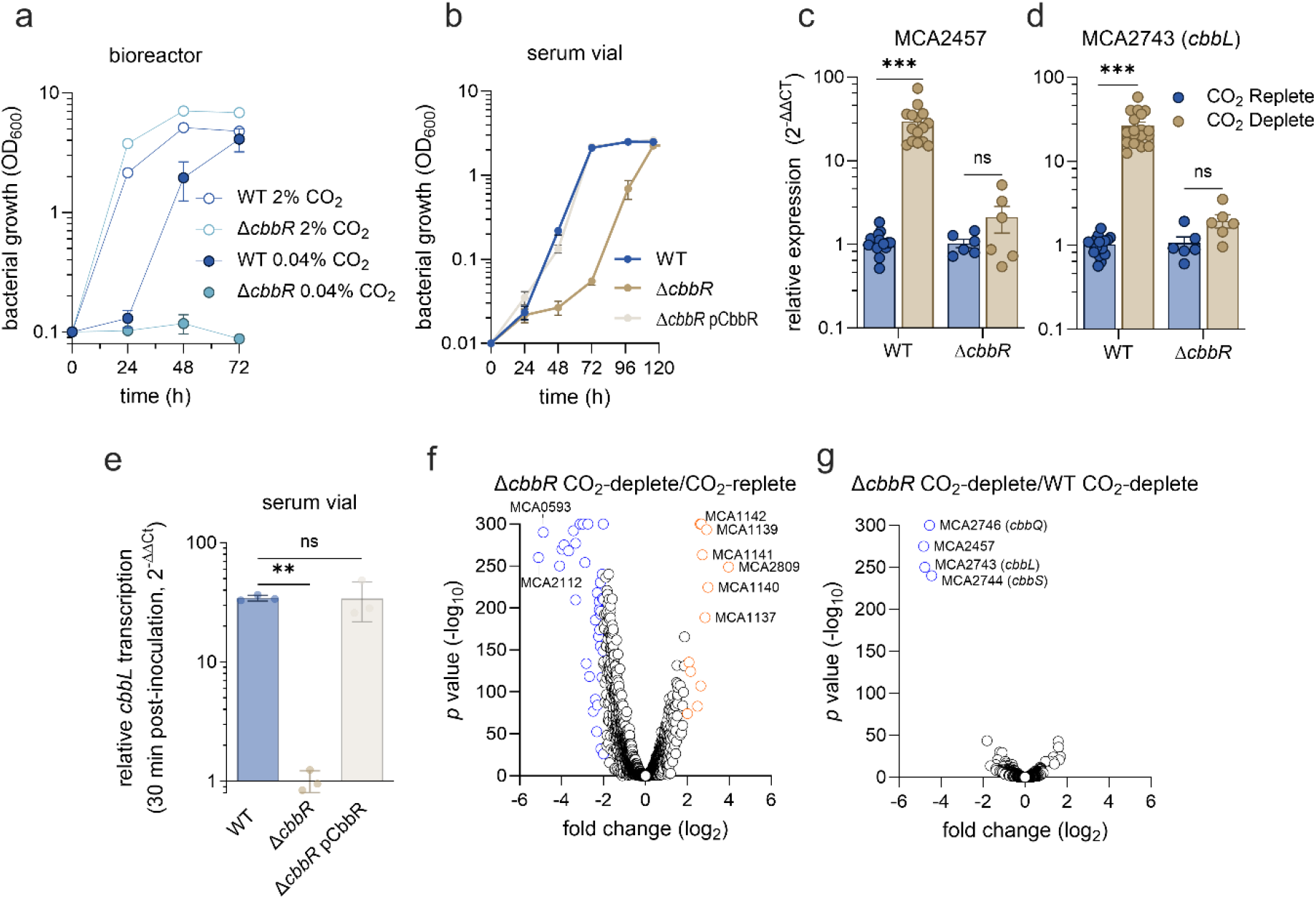
CbbR activates RubisCO transcription in response to CO_2_ starvation in *M. capsulatus.* a) Wildtype (WT) and CbbR-deficient (Δ*cbbR*) *M. capsulatus* cultivated in a bioreactor with continuous gas sparging of 20% CH_4_/2% CO_2_ in air (empty symbols) or 20% CH_4_/0.04% CO_2_ in air (filled symbols). b) Growth of WT, Δ*cbbR,* and the complementing Δ*cbbR* pCbbR strain in sealed serum vials supplied with 20% CH_4_ in air. c-d) RT-qPCR analysis of (c) MCA2457 and (d) MCA2743/*cbbL* gene expression in WT and Δ*cbbR* bioreactor cultures either unstarved (blue bars) or CO_2_ starved for 1 h. e) RT-qPCR analysis of *cbbL* transcription in WT, Δ*cbbR,* and the complementing Δ*cbbR* pCbbR strain 30 min post-inoculation into sealed serum vials with 20% CH_4_ in air (no CO_2_ supplementation) compared to vials with 20% CH_4_/8% CO_2_ in air. f) Volcano plot of differentially expressed genes in the Δ*cbbR* strain 1 h after CO_2_ removal from the bioreactor sparge gas stream (CO_2_-deplete) compared to unstarved controls. g) Volcano plot of differentially expressed genes between Δ*cbbR* and WT *M. capsulatus* 1 h after CO_2_ removal from the bioreactor sparge gas stream (CO_2_-deplete). Data in a-e represent the mean ± SEM (n=3-8) from at least two independent experiments, and data in f-g are based on RNA sequencing from 3 biological replicates for each condition. \*\**p* ≤ 0.01, *** *p* ≤ 0.001 from unpaired *t*-tests. ns, not statistically significant.

Next, we evaluated whether CbbR regulates RubisCO and MCA2457 transcription via direct binding to the gene promoters. Electromobility shift assays using purified CbbR-6xHis and a biotinylated amplicon containing the *cbbL* or MCA2457 promoter elements (P*_cbbL_* or P_MCA2457_), respectively, supported that CbbR directly binds to both P*_cbbL_* and P_MCA2457_ promoter regions (Fig. 6a), consistent with the RT-qPCR analysis in the Δ*cbbR*::Kn^R^ knockout strain (Fig. 5). LTTRs are often allosterically regulated by metabolite effectors that can alter transcription factor-DNA interactions (39, 44–47). We performed untargeted metabolomics comparing the intracellular concentration of metabolites under CO_2_-deplete and unstarved conditions to identify candidates that may be involved in CbbR allosteric regulation and activation of the CO_2_ starvation response (Table S15). Several metabolites were significantly altered in cells within 5 h following CO_2_ starvation (Fig. 6b). Notably, the RubisCO substrate, RuBP, and the product of RubisCO-dependent RuBP oxygenation, 2-phosphoglycolate (2PG) were significantly elevated in CO_2_-deplete cells compared to unstarved cells (RuBP, log_2_FC = 3.91; 2PG, log_2_FC = 2.83) Several tricarboxylic acid cycle metabolites were also altered in response to CO_2_ starvation, including 2-ketoglutarate (log_2_FC = 2.16), fumarate (log_2_FC = -0.97), and malate (log_2_FC = -2.03). The amino acids aspartate, ornithine, and glutamine were significantly decreased (log_2_FC = -4.48, -2.05, -0.70, respectively). Collectively, the significant decreases in these amino acids, 2-ketogluterate, and the transcriptional repression of nitrogen uptake and translational machinery indicate that nitrogen assimilation and metabolism is dysregulated in CO_2_-starved *M. capsulatus*.

**Figure 6.**
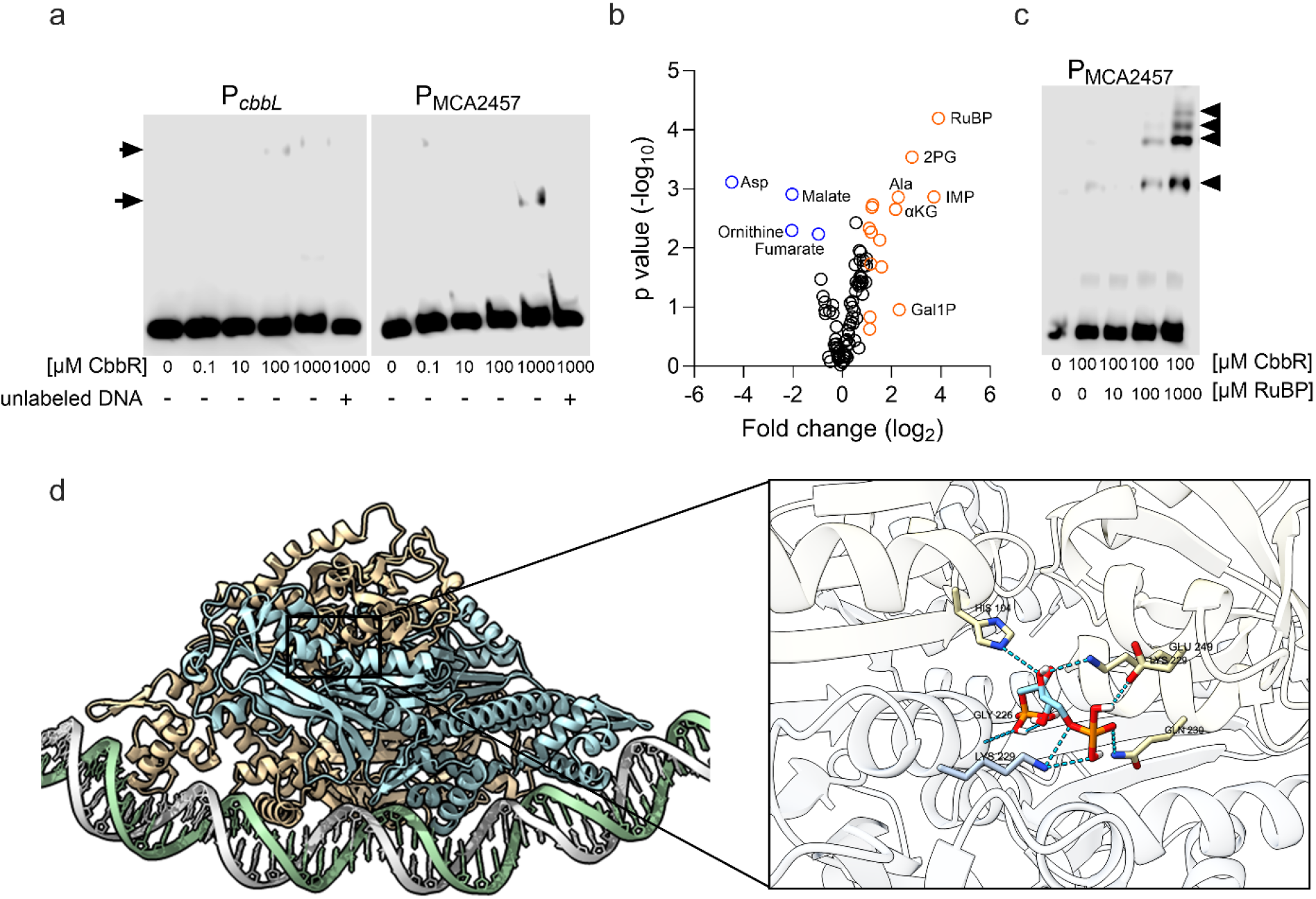
CbbR is allosterically regulated by ribulose-1,5-bisphosphate to directly activate carbon capture gene promoters during CO_2_ limitation. a) Electrophoretic mobility shift assays evaluating CbbR-6xHis binding to biotinylated P_MCA2743_ (*cbbL*) and P_MCA2457_ promoter regions. Arrows indicate CbbR-promoter DNA complexes. b) Differential abundance of intracellular metabolites in CO_2_ starved vs. unstarved *M. capsulatus* cells. c) CbbR-6xHis binding to the P_MCA2457_ promoter with increasing concentration of ribulose-1,5-bisphosphate (RuBP). Arrows indicate CbbR-P_MCA2457_ complexes. d) Structural model of an RuBP-activated CbbR tetramer bound to the P_MCA2457_ promoter region. CbbR dimers are colored in blue and tan. The putative RuBP binding site residues between CbbR dimers are shown (inset).

Given that 1) RuBP and 2PG were elevated in cells eliciting the CO_2_ starvation response and 2) both RuBP and 2PG are known effectors of CbbR homologs in *Synechocystis* sp. PCC 6803 and *Rhodobacter sphaeroides* (39, 48), we evaluated the effect of these putative effectors on CbbR binding to P*_cbbL_* and P_MCA2457_ promoters. RuBP enhanced CbbR binding to both P*_cbbL_* and P_MCA2457_ with multiple shifted CbbR-DNA bands observed at higher RuBP concentrations (Fig. 6c and Fig. S4), which may be due to various RuBP-induced CbbR oligomeric states as previously hypothesized for *Rhodobacter sphaeroides* CbbR (49). In contrast, 2PG had no effect on CbbR binding to either promoter (Fig. S4). Boltz-2 (50) was used to predict the residues coordinating RuBP within effector-bound CbbR and its interactions with the P_MCA2457_ promoter (Fig. 6d). The predicted structure (ipTM = 0.5673) indicates that CbbR may bind to DNA as a tetramer (Fig. 6d). Taken together, these results support that CbbR is allosterically modulated by the RubisCO substrate, RuBP, to function as a transcriptional activator of RubisCO and MCA2457 in response to CO_2_ limitation.

## Discussion

Methanotrophs play a fundamental role in global biogeochemical cycles and are important members of terrestrial and aquatic microbial communities, so understanding how they respond to changing environmental conditions is of interest. As specialists with the ability to convert CH_4_ to multi-carbon compounds, methanotrophs are also promising microbial chassis for biotechnology applications targeting atmospheric CH_4_ mitigation or conversion of CH_4_-rich gas streams. Our investigations identify a coordinated transcriptional response to CO_2_ limitation in the RubisCO-encoding methanotroph, *Methylococcus capsulatus*, that includes the induction of inorganic carbon metabolic enzymes and the concomitant repression of nitrogen metabolism and translational machinery. This global transcriptional response is comparable to starvation responses characterized in other microorganisms where nutrient limitation triggers a coordinated reallocation of cellular resources toward acquisition of the limiting nutrient (32, 33, 51). We show a *M. capsulatus* CbbR homolog is a central regulator of this response, further expanding the role of CbbR-family transcriptional regulators in methylotrophs beyond their canonical roles in regulating inorganic carbon fixation in autotrophs. The CO_2_ starvation response identified here is consistent with the physiological requirement of CO_2_ for *M. capsulatus* growth (19) and underscores a metabolic link between inorganic and organic carbon and nitrogen metabolisms in this methanotroph.

Although *M. capsulatus* derives most of its carbon and energy from CH_4_, our results support CO_2_ is an important component of this bacterium’s physiology. Early seminal studies demonstrated that *M. capsulatus* assimilates CO_2_ during growth on CH_4_ and provided evidence for a functional CBB cycle (26, 52). More recently, genetic and physiological analyses showed that RubisCO and CO_2_ are necessary for growth of *M. capsulatus* Bath (19), establishing inorganic carbon as an essential component of the organism’s metabolism rather than simply a metabolic by-product of CH_4_ oxidation. Our observation that *M. capsulatus* can be cultivated at or slightly above atmospheric CO_2_ concentrations (0.04%) supports that this organism has sufficient capacity for inorganic carbon acquisition under environmentally relevant CO_2_ concentrations. Indeed, the CO_2_ starvation transcriptional response not only included the induction of RubisCO but also carbonic anhydrases and bicarbonate transporters, probable components of a carbon concentrating mechanism that would enhance CO_2_ assimilation by *M. capsulatus*. The up-regulation of carbon concentration machinery along with RubisCO is a common strategy employed by autotrophic bacteria to persist in low CO_2_ environments (53–57), and potential interactions between these proteins to mediate *M. capsulatus* inorganic carbon assimilation warrants further investigation.

The cessation of *M. capsulatus* growth at sub-atmospheric CO_2_ concentrations is linked to a significant decrease in transcription of nitrogen metabolism and translational machinery genes, a response consistent with the conserved (p)ppGpp-dependent stringent response to nutrient limitation characterized in an array of bacteria (58–61). Notably, *M. capsulatus* encodes two RelA/SpoT (p)ppGpp synthetase homologs (MCA1920 and MCA2023), but (p)ppGpp biosynthesis has not previously been evaluated in this bacterium or other methanotrophs. The overlap between the CO_2_ starvation response found here and stringent response-associated genes suggests that CO_2_ limitation may be integrated into a broader physiological transition from growth to resource conservation in *M. capsulatus*. In this model, CbbR promotes the acquisition and utilization of scarce inorganic carbon to maintain carbon homeostasis and survive CO_2_ limitation, while the stringent response simultaneously suppresses energetically expensive biosynthetic processes to limit growth. Defining the regulatory interactions between the CO_2_ starvation and stringent responses will be an important next step toward understanding carbon and nitrogen balancing in C1-utilizing microbes and enhancing growth of these bacteria for biotechnology applications.

In several chemolithoautotrophic and phototrophic bacteria, CbbR is required for expression of the CO_2_-fixation machinery in response to CO_2_ limitation, positively regulating transcription upon allosteric activation by elevated CBB cycle metabolite pools or NADPH (39, 41, 42, 47, 62). Similarly, the *M. capsulatus* CbbR is allosterically regulated by RuBP, which spikes in the absence of CO_2_. Although the *M. capsulatus* CbbR positively regulates several gene targets in response to CO_2_ limitation, its regulon appears to be limited to the RubisCO operon and the MCA2457 gene encoding a putative carbonic anhydrase. Notably, several genes were still induced under CO_2_-deplete conditions in the CbbR-deficient Δ*cbbR* strain, suggesting additional regulators may be involved in the *M. capsulatus* CO_2_ starvation response. Repression of translational machinery and nitrogen metabolism genes also appear to be independent of CbbR and likely represent a more generalized stringent response to carbon starvation rather than inorganic carbon specifically. Indeed, a similar stringent transcriptional response was observed in the related gammaproteobacterial methanotroph *Methylotuvimicrobium buryatense* 5GB1C during cultivation with low (500 ppm) CH_4_ availability (63).

CbbR homologs, termed QscR, have previously been identified in methylotrophic bacteria where they are critical regulators of C1 metabolism (64–66). In the facultative methylotroph *Methylobacterium extorquens*, QscR functions as a positive regulator of the serine cycle, the principal C1-assimilation pathway in alphaproteobacterial methylotrophs and methanotrophs (64, 66). Loss of QscR severely compromises *M. extorquens* growth on C1 substrates and reduces expression of multiple serine-cycle genes (65). Similar to our findings, the *M. extorquens* QscR is allosterically regulated by RuBP (66) which is synthesized by a phosphoribulokinase encoded by *M. extorquens,* although this bacterium does not encode RubisCO (67). It seems the regulatory role of CbbR-family proteins has expanded from exclusively regulating inorganic carbon acquisition and metabolism in autotrophs to also coordinating metabolic pathways responsible for assimilation of reduced C1 compounds in methylotrophs. The parallels of the CO_2_ starvation response identified here with known autotrophic responses to CO_2_ availability highlight an evolutionary relationship between autotrophy and methanotrophy. Our results support that CbbR/QscR represents an evolutionary link between autotrophic and methylotrophic metabolisms, which have distinct but overlapping C1 biochemical pathways. Notably, *M. capsulatus* inorganic carbon metabolism is not independent from C1 metabolism because CO_2_ represents both an essential metabolic substrate and a product of reduced C1 substrate oxidation. As such, the development of a regulatory system capable of sensing CO_2_ availability provides an effective mechanism to coordinate inorganic carbon availability with the energetic and biosynthetic demands of C1 metabolism.

Taken together, our investigations establish CbbR as an important component of CO_2_ homeostasis in *M. capsulatus* and provide a framework for understanding how inorganic carbon-responsive regulation has been integrated into methanotrophic physiology. The coordinated induction of CO_2_ scavenging machinery and a stringent starvation response by *M. capsulatus* support that CO_2_ limitation is not only perceived as a substrate deficiency but also as a broader physiological stress requiring coordinated metabolic reprogramming. This CO_2_ starvation transcriptional response may represent a general strategy that enables methanotrophs to persist in environments with fluctuating CH_4_ or CO_2_ concentrations. Notably, the low CO_2_-induced genes/enzymes that facilitate efficient CO_2_ assimilation by *M. capsulatus* identified in this study represent rational engineering targets to potentially improve carbon assimilation by these microbes and advance their use in C1 conversion biotechnologies.

## Methods

### Bacterial strains and cultivation

Bacterial strains and plasmids used in this study are listed in Tables S16 and S17, respectively. *E. coli* strains were cultivated using lysogeny broth (Lennox) solid or liquid medium containing 100 µg/mL ampicillin, 50 µg/mL spectinomycin, 50 µg/mL kanamycin, or 15 µg/mL gentamycin as necessary. *M. capsulatus* Bath strains were maintained on nitrate mineral salts (NMS) solid medium in sealed chambers containing 20% CH_4_ and 2% CO_2_ at 37°C as previously described (19). *M. capsulatus* strains streaked from a -80°C stock (cryopreserved in 8% DMSO in NMS) were passaged weekly for a maximum of four weeks before being replaced. Plate biomass was used to inoculate 10mL of NMS medium in a 150 mL sealed serum vial or 20 mL of NMS medium in a 250 mL vial to OD_600_ = 0.01 as previously described (19). 50 µg/mL kanamycin, 30 µg/mL gentamicin, 50 µg/mL spectinomycin, or 50 µg/mL erythromycin were supplemented to liquid and solid NMS medium as necessary. *M. capsulatus* Bath strains were also cultivated in a custom mid-throughput gas bioreactor with continuous gas flow as previously described (19). Briefly, 150 mL Kimax tubes containing 100 mL of NMS with 75 µg/mL cycloheximide and selective antibiotic, if applicable, were inoculated at OD_600_ = 0.1 (unless otherwise stated) using a seed culture grown overnight to mid-log phase (OD_600_ ∼ 0.5-1) in a serum vial. CH_4_, CO_2_, and air were mixed with mass flow controllers (Alicat) and supplied to cultures at 1 volume gas/volume culture medium/min (vvm) through a stainless-steel sparge stone. Where appropriate, culture viability was determined by quantifying viable cells as colony forming units through serial dilutions followed by plating on solid media.

### Gas chromatography

CH_4_ consumption and CO_2_ evolution were measured using a SRI 8610C gas chromatograph (GC) equipped with a 6’ x 1/8in Hayesep D packed column and thermal conductivity and flame ionizing detectors using helium and hydrogen as carrier gases. For serum vial cultivations, 0.5 mL of vial headspace was removed and injected directly into the GC on-column injector using a Heinke-ject Air Tite 1mL syringe. To quantify CH_4_ and CO_2_ in the effluent stream from the mid-throughput bioreactor, samples were taken directly from the vial exhaust port using 1/8 in Nalgene tubing. CH_4_ and CO_2_ were quantified via regression analysis compared to known standards.

### ATP and NADP(H) quantification

Bioreactor cultures inoculated at OD_600_ = 0.1 were supplied with 20% CH_4_/2% CO_2_ in air for 3 h (unstarved) after which CO_2_ was removed from the gas stream (CO_2_-deplete). Following 3 h of CO_2_ depletion, cells were harvested for quantification of NADP(H) and ATP. NADPH and NADP^+^ were quantified using a modified enzymatic MTT cycling assay (68). Briefly, CO_2_-starved cells were pelleted at 8000 x g for 5 min at 4°C and resuspended in 50 µL 0.2 M NaOH (NADPH) or 0.2 M HCl (NADP^+^) and incubated at 55°C for 10 min. Samples were then transferred to ice for 10 min before being neutralized with an equivalent volume of 0.2 M HCl (NADPH) or 0.2 M NaOH (NADP^+^) added dropwise while gently vortexing. 100 µL reactions containing 10 µL of clarified cell lysate, 45 µL reaction master mix (1 M bicine, 12.7 mM glucose-6-phosphate, 4.2 mM MTT, and 16 mM phenazine ethanosulfate premixed in a 1/1/1/0.5 ratio), 1 µL of 1 mg/mL glucose-6-phosphate dehydrogenase, and 44 µL nuclease free H_2_O were prepared in a 96-well plate. The reduction of MTT to formazan was measured spectrophotometrically at A_540_ at 30 sec intervals for 4 min using a Biotek SynergyMx microtiter plate reader. The slope was used to quantify NADPH and NADP^+^ via linear regression compared to known NADP(H) standards.

ATP was quantified using the ATP Detection Assay Kit (Cayman Chemicals) following the manufacturer’s protocol. 100 mL cultures of unstarved and CO_2_-starved cells 3 h after CO_2_-depletion were used to generate ATP-containing lysates. ATP-dependent luciferase luminescence was measured using a Biotek SynergyMx microtiter plate reader and ATP quantified compared to known standards supplied with the assay kit.

### RNA extraction, RNA sequencing, and quantitative reverse transcription PCR (RT-qPCR)

*M. capsulatus* seed cultures were inoculated from plate biomass and grown overnight with 20% CH_4_ in air at 37°C in serum vials to OD_600_ ∼ 1. Serum vial seed cultures were subcultured to OD_600_ = 0.1 in NMS to a total volume of 100 mL. For CO_2_-depletion experiments, 20% CH_4_/2% CO_2_ in air was sparged at 1 vvm for 3 h followed by removal of the CO_2_ from the gas mixture. For CO_2_-replete experiments, 20% CH_4_ in air was sparged at 1 vvm for 3 h followed by the addition of 2% CO_2_ to the gas mixture. At the indicated time, 40 mL of culture was immediately added to a 50 mL conical tube in an ice bucket containing 10 mL ice-cold stop solution (5% phenol/95% ethanol) and incubated for 20 min on ice to stabilize RNA. Fixed cells were pelleted for 10 min at 6000 x g, 4°C and used immediately for RNA extraction or stored at -20°C. RNA was extracted using the High Pure RNA Isolation Kit (Roche) following the manufacturer’s instructions, which included a 10 min lysozyme incubation and an on-column DNase treatment. Any remaining residual DNA was removed with Turbo DNase (ThermoFisher Scientific) followed by clean-up with a Monarch Spin RNA Cleanup Kit (New England Biolabs) to obtain high-quality RNA. RNA quality was determined by gel electrophoresis and Agilent TapeStation prior to RNA sequencing or RT-qPCR. Sequencing libraries were generated using the Illumina Stranded Total RNA Prep with Ribo-Zero Plus rRNA Depletion Kit and sequenced with an Illumina NextSeq 500 system at the University of North Texas Genomics Center. Sequence reads with ≥ Q30 scores (>10 million/sample) were processed using the BBDuk plugin and aligned to the *M. capsulatus* Bath genome (NC_002977) using Geneious software. Expression levels were calculated after mapping and compared between unstarved, CO_2_-deplete, or CO_2_-replete conditions to determine differentially expressed genes with the Geneious DESeq2 plugin. Illumina sequence data have been deposited in the NCBI BioProject Database under accession number PRJNA1548772.

For RT-qPCR, cDNA was synthesized from 1 µg purified total RNA using the SuperScript IV Vilo Master Mix following the manufacturer’s protocol (ThermoFisher Scientific). 1 µL cDNA was used as template in a 10 µL reaction with PowerUp SYBR Green Master Mix (Applied Biosystems) and primers (Table S18) designed using the PrimerQuest Tool (Integrated DNA Technologies). Target gene amplification was performed using a Quantstudio 3 Real-Time PCR System (Applied Biosystems) with standard amplification parameters. Gene expression was determined by normalizing the target cycle threshold (Ct) to that of the housekeeping *rpoD* gene encoding the RNA polymerase σ70. Relative gene expression between conditions was calculated using the ΔΔCt method. For graphing purposes, genes with *p* = 0 were assigned the minimum *p*-value > 0 from the related data set. Actual *p*-values can be found in the supplementary data tables.

### CbbL epitope tagging and western blotting

The pCBBL3xFLAG construct for ectopic expression of RubisCO (CbbLS) with a 3xFLAG epitope tag on the CbbL N-terminus was generated by isothermal assembly of two linear fragments using NEBuilder 2x HiFi DNA Assembly Master Mix (New England Biolabs). The two fragments were amplified using Q5 High-Fidelity 2X Master Mix (New England Biolabs): 1) the pLSQ backbone (37) linearized by PCR using primers oCAH1751 and oCAH17, and 2) a *cbbL* ORF with an N-terminal 3xFLAG tag sequence amplified from a synthetic g-block DNA fragment (3XFlag-*cbbLS*) synthesized by Integrated DNA Technologies using primers oCAH1629 and oCAH1760 (Table S18). The isothermal assembly reaction was transformed into DH10B *E. coli* and transformants selected on LB with kanamycin. The resulting plasmid, pCBBL3xFLAG, sequence was confirmed via whole plasmid sequencing (Genewiz) and subsequently transferred to *M. capsulatus* Bath via biparental mating using S17-1λpir *E. coli* as the donor strain. Kanamycin-resistant transconjugants were confirmed to harbor the pCBBL3xFLAG plasmid via PCR and whole-plasmid sequencing of isolated plasmid DNA.

Seed cultures of the pCBBL3xFLAG *M. capsulatus* strain grown overnight in serum vials were sub-cultured to OD_600_ = 0.1 in 100 mL of NMS medium in the mid-throughput gas bioreactor and cultivated with continuous supply of 20% CH_4_/2% CO_2_ in air for 3 hours followed by CO_2_ removal from the gas stream. At the indicated time post CO_2_ starvation, 10 mL culture was pelleted by centrifugation at 5000 x g at 4°C for 10 min. Cell pellets were resuspended in 200 µL B-PER reagent (ThermoFisher Scientific) for protein extraction. The protein concentration in whole-cell lysates was determined using the Pierce 660nm Protein Assay Reagent (ThermoFisher Scientific) compared to BSA standards. Lysates were normalized based on protein concentration and then equally mixed with 2x Laemmli buffer containing 100 mM dithiothreitol and heated at 85°C for 5 min. 5 µg of protein lysate was resolved on a 12% SDS PAGE gel before being transferred to a PVDF membrane using an XCell SureLock MiniCell and Xcell II Blot Module (ThermoFisher Scientific). Prior to blocking, membranes were stained with 0.1% w/v Ponceau S in 5% acetic acid to confirm equivalent protein loading. Following Ponceau S staining, membranes were washed with ddH_2_O and blocked for 1 h in 5% (w/v) non-fat milk in TBST (blocking buffer) at room temperature. The membrane was then probed with the M2 anti-FLAG antibody (Sigma Aldrich) diluted 1:1000 in blocking buffer overnight at 4°C. The membrane was washed 5x by rocking 10 min with TBST and then incubated for 1 h with an anti-mouse-HRP conjugated secondary antibody (Santa Cruz Biotechnology) diluted 1:20,000 in blocking buffer. The membrane was then washed 5x by rocking 10 min with TBST. HRP chemiluminescence was detected using the SuperSignal West Femto substrate (ThermoFisher Scientific) and imaged with a C-DiGit Blot Scanner (LICOR). Band intensity was quantified using ImageJ V2.14.0 as previously described (69).

### 13CO2 isotopic labeling and LC-MS/MS

To determine the incorporation of CO_2_ into *M. capsulatus* central metabolites, 10 mL of culture from bioreactor cultivations supplied with 20% CH_4_/2% CO_2_ (unstarved) or CO_2_-starved for 3 h were transferred to sealed serum vials with atmospheres containing 20% ^12^CH_4_ and 8% ^13^CO_2_ and incubated for 5 min at 37°C with shaking. After incubation, ^13^C-labeled cells were isolated by vacuum filtration using a Buchner funnel equipped with a 0.2 µm nylon membrane, which was quickly transferred to a conical tube and frozen in liquid nitrogen. The membranes were freeze-dried for 48 h and metabolites were then extracted from dried cells using a boiling water extraction method as previously described (70). Phosphorylated central metabolites and amino acids were separated and quantified via LC-MS/MS at the UNT Bioanalytical Facility as previously described (71). Total quantity of ^13^C-labeled metabolites was calculated by multiplying the quantity of metabolite determined by quantitative targeted metabolomics (see below) by the isotopic ratio determined in the ^13^CO_2_ labeling experiment.

### CRISPRi library construction and Oxford Nanopore Technologies sequencing

Synbio Technologies designed and synthesized a 6,220 sgRNA oligo pool which included 15 sgRNAs per the 388 DEGs identified in response to CO_2_ starvation plus 400 non-targeting sgRNAs with randomly generated spacers. The sgRNA oligos (120 bp) possess 5’ and 3’homologous sequence for cloning into the p01sgRNA plasmid (37). The pooled sgRNA library was amplified via PCR using primers oCAH1416/1417 and Q5 polymerase (New England Biolabs) following the manufacturers recommended cycling parameters for 20 cycles. The amplicon was assembled with p01sgRNA amplified with oCAH1190/oCAH119 via isothermal assembly and transformed into high-efficiency DH10B competent cells (New England Biolabs) as previously described (37). The sgRNA library was isolated from pooled DH10B transformants (∼10^6^ CFU/mL) and then transformed into S17-1λpir *E. coli* for conjugative transfer to an *M. capsulatus* Bath strain with a chromosomally integrated P_tet_-dCas9 to generate the targeted CRISPRi library. The methanotroph targeted CRISPRi library size was quantified via serial dilution plating an aliquot of the mating prior to plating the remaining mating biomass on selective medium. A bacterial lawn of ∼1 x 10^6^ individual transconjugant CFUs were pooled and aliquoted for long term -80°C storage and genomic DNA isolated to determine library coverage. For screening, the targeted sgRNA library was cultivated in liquid NMS overnight and subsequently subcultured to OD_600_ = 0.01 in fresh NMS medium with (CRISPRi induction) or without 0.2 µg/mL anhydrotetracycline inducer. After 72 h of selection, genomic DNA was extracted from the cells using a Zymo DNA extraction kit following the manufacturer’s protocol. 50 ng gDNA was used as a template to amplify a ∼250 bp amplicon containing the sgRNA sequence with Q5 polymerase using primers oCAH1416/oCAH1417 and the same reaction conditions as above (20 cycles). The amplicons from uninduced and induced conditions were barcoded with the native barcoding kit 24 V14 following the manufacturers protocol (Oxford Nanopore Technologies, ONT) and sequenced using a MinION sequencing device. sgRNA target sequences were extracted from the sequence data and mapped to the *M. capsulatus* genome to calculate sgRNA relative fitness between induced and uninduced samples using Geneious software following previously described bioinformatic workflows (37).

### M. capsulatus ΔcbbR knockout strain construction

The MCA3039 KO (Δ*cbbR::kn)* strain was created through allelic exchange mutagenesis as previously described (19). Homologous regions 1 kbp upstream and downstream of the MCA3039 open-reading-frame were amplified from *M. capsulatus* genomic DNA using primers oCAH579/oCAH580 and oCAH581/oCAH582, respectively. An FRT-flanked kanamycin resistance cassette with homology to the upstream and downstream flanks was amplified from pKD13 using primers oCAH482/oCAH487. The three fragments were assembled with the pK18mobpheS methanotroph suicide plasmid backbone linearized via PCR with primers oCAH576/oCA577. The pK18*ΔcbbR* plasmid was transferred to *M. capsulatus* Bath via conjugation and kanamycin-and 4-chlorophenylalanine-resistant transconjugants were selected on NMS medium. Primers oCAH579/oCAH582 flanking the MCA3039 locus were used to verify replacement of the MCA3039 ORF with the kanamycin cassette.

The pCbbR plasmid for complementation was constructed by amplifying a fragment containing the MCA3039 ORF with its native promoter from *M. capsulatus* gDNA using primers oCAH1330/oCAH1331 which was assembled with the pBBR1-based replicative plasmid pDS1 amplified with oCAH16/oCAH17, replacing the pDS1 promoter and mRFP gene. The sequence-confirmed plasmid was then moved to S17-1λpir *E. coli* for transfer to the Δ*cbbR::kn* strain via conjugation. Kanamycin- and gentamicin-resistant transconjugants were verified to harbor the pCBBR plasmid and lack the MCA3039 ORF via PCR.

### CbbR purification

The *cbbR* ORF amplified with primers oCAH1733/oCAH1734 was cloned into pET22b linearized with primers oCAH1579/oCAH1580 via isothermal assembly. The sequence-confirmed plasmid was transformed into chemically competent BL21 Star (DE3) *E. coli* (ThermoFisher Scientific) for expression and purification of a C-terminal 6xHis-tagged CbbR variant.

Initial attempts to purify CbbR resulted in negligible quantities of soluble protein. Previous studies have shown that co-expression of a GroEL family chaperone can increase CbbR solubility (72). Thus, we identified an *M. capsulatus* GroEL homolog (encoded by MCA0707) and cloned the gene with its native ribosome binding site downstream of the *cbbR*-6xHis to generate an IPTG-inducible polycistronic operon (pSAL01). The construct was sequence-confirmed by whole plasmid sequencing (Genewiz) and then transformed into BL21 Star (DE3) *E. coli*.

For CbbR-6xHis and GroEL coexpression, BL21 Star (DE3) *E. coli* harboring pSAL01 was cultured overnight and then subcultured 1:100 in 1 L fresh LB medium containing ampicillin in a 3 L Fernbach flask and incubated at 37°C with shaking at 200 rpm for ∼ 3 h when the culture reached OD_600_ ∼ 0.5. The culture was then induced with 1 mM IPTG for 16 h at 16°C with shaking at 200 rpm. After induction, the culture was transferred to 0.5 L centrifuge tubes and placed on ice for 10 minutes.

The cells were centrifuged at 5000 x g at 4°C for 10 min and the cell pellets were resuspended in 50 mL of lysis buffer (50 mM Tris-HCl pH 7.5, 100 mM NaCl, 10 mM imidazole, cOmplete EDTA free protease inhibitor (Roche) per liter of culture and frozen at - 80°C. The resuspended pellets were subjected to three freeze thaw cycles followed by incubation with 10 µL of lysozyme (50 µg/µL) for 20 min at 37°C. Cells were then placed on ice and sonicated five rounds of two-minute increments using a Fisher FB120 sonicator and a 0.63 cm probe, set at 40% amplitude with an alternating pulse time of 5 sec and a rest time of 5 sec for a total of 10 min of sonication. Lysates were then centrifuged at 20,000 x g at 4°C for 1 h to isolate the soluble fraction.

CbbR-6xHis was purified by affinity chromatography using a gravity flow column packed with 1 mL bed volume of HisPur Ni-NTA Resin (ThermoFisher Scientific). Clarified lysate was passed through the column twice and then the resin was washed with 20 mL of wash buffer (100 mM Tris-HCl, 150 mM NaCl, and 20 mM imidazole). CbbR-6xHis was eluted from the column by adding 4 mL of elution buffer (100 mM Tris-HCl, 150 mM NaCl and 300 mM imidazole). The eluate was then passed through a Superdex 200 10/300 column using an AKTA Pure FPLC with a carrier buffer consisting of 100 mM Tris-HCl pH 8.0 and 150 mM NaCl.

Protein was detected using a U9-L UV detector at 280 nm and fractions corresponding to the correct molecular weight were collected, pooled, and concentrated with a 10 kD MWCO Amicon Ultra Centrifugal Filter. Column flow through, wash, and eluate samples were evaluated by SDS PAGE.

### Electromobility shift assays

Electromobility shift assays (EMSA) were performed using the Lightshift Chemiluminescent EMSA assay kit (ThermoFisher Scientific) following the manufacturer protocol. 150 bp promoter regions directly 5’ of the start codon for MCA2457 and *cbbL* were amplified with biotinylated forward primers to generate labeled DNA probes (oCAH1510/oCAH1511 for P*_MCA2457_* and oCAH1512/oCAH1513 for P*_cbbL_*) or unmodified forward primers to generate specific competitor probes (oCAH1515 for P*_MCA2457_* and oCAH1516 for P*_cbbL_*). Transcription factor-DNA binding reactions (20 µL) were incubated for 1 h at room temperature and contained increasing concentrations of CbbR-6xHis as indicated, 20 fmol of biotinylated probe DNA, 0.05% NP-40, 1 mM DTT, 2.5% glycerol, and 1 µg Poly dI-dC oligonucleotides (ThermoFisher Scientific). For direct competition reactions, 4 pmol of unlabeled probe was added to the reactions. To determine the effect of metabolites on DNA binding, ribulose-1,5-bisphosphate or 2-phosphoglycolate were added to the binding reactions at indicated concentrations.

Protein-DNA binding reactions were loaded onto a 6% 0.5x Tris-Borate-EDTA (TBE) gels (pre-run for 30 mins at 100 V prior to sample loading) and electrophoresed at 120 V for 75 min. Protein and DNA were then transferred to a Novex Nylon+ 0.45 µm pore membrane on ice in 0.5x TBE for 30 min at 380 mA. After transfer, the membrane was fixed with a UV Stratalinker 1800 (Stratagene) using the auto-crosslink function. Membrane blocking, washing, labeling and detection were performed following the manufacturer’s instructions using the provided kit reagents. Blots were visualized on a C-digit blot scanner using Imagestudio V6.0 (Licor).

### Targeted metabolomics

Serum vial seed cultures were cultivated for ∼16 h, diluted to OD_600_ = 0.1 in 100 mL NMS in 150 mL kimax bioreactor tubes, and incubated in the mid-throughput bioreactor at 37°C with continuous sparging of 20% CH_4_/2% CO_2_ in air for 3 h (unstarved). After 3 h, CO_2_ was removed from the sparge gas and 20% CH_4_ in air was supplied for 5 h (CO_2_-deplete). Unstarved and CO_2_-deplete cells (100 mL of OD_600_ = 0.1) were pelleted at 5000 x g for 10 min at 4°C, and the cell pellet was immediately frozen with liquid nitrogen to quench metabolism. Cells were lyophilized for 48 h at -80° C using a Freezone 12-liter console freeze dryer (Labconco). Lyophilized samples (∼4 mg) were resuspended in 1 mL ddH_2_O and metabolites were extracted by boiling as previously described (73). Metabolites were then separated and quantified with a 1290 Infiniti II ultra-high-performance liquid chromatograph (Agilent) coupled to an AB QTRAP 6500+ mass spectrometer (SCIEX) system as previously described (71).

### CbbR-promoter DNA complex modeling

A complex of tetrameric CbbR, the RubisCO *cbbLSQ* operon promoter region (150 bp directly up-stream of the *cbbL* ORF) and two molecules of RuBP was modeled using Boltz-2 (50). Affinity for and position of RuBP in the CbbR effector binding domain was determined using Autodock vina v1.2.0. Complex visualization and RuBP docking were rendered in ChimeraX v1.10.

### Illustrations

Data were graphed using GraphPad Prism 10 software. Circos plots were generated using Circa (https://circa.omgenomics.com/). Workflow schematics were created with BioRender.com.

### Statistical analysis

Statistical analysis was performed using GraphPad Prism 10 software. The data between two groups were analyzed using unpaired t-tests. The determination of statistical significance between multiple comparisons was achieved using analysis of variance (ANOVA) followed by Dunnett’s multiple comparisons test. Data were considered statistically significant when P ≤ 0.05.

## Acknowledgments

We would like to thank Dr. Danyang Shao for computational support with sgRNA spacer extraction, Robyn Alba for assistance with pK18*ΔcbbR* cloning, and Liam Jouvenat for RT-qPCR technical assistance.

## Funding

BioDiscovery Institute Seed Grant (CH)

National Science Foundation MCB award # 2225776 (CH)

National Science Foundation MCB CAREER award #2441696 (CH)

## Author contributions

Conceptualization: SL, CH

Formal analysis: SL, JH, YCY, CH

Funding acquisition: CH

Investigation: SL, JH, YCY, CH

Methodology: SL, JH, CH

Supervision: CH

Visualization: SL, CH

Writing—original draft: SL, CH

Writing—review & editing: SL, JH, YCY, CH

## Competing interests

Authors declare that they have no competing interests.

## Data and materials availability

All data are available in the main text or the supplementary materials. Illumina sequence data have been deposited in the NCBI BioProject Database under accession number PRJNA1548772.

## Supplementary Figures

**Fig. S1.**
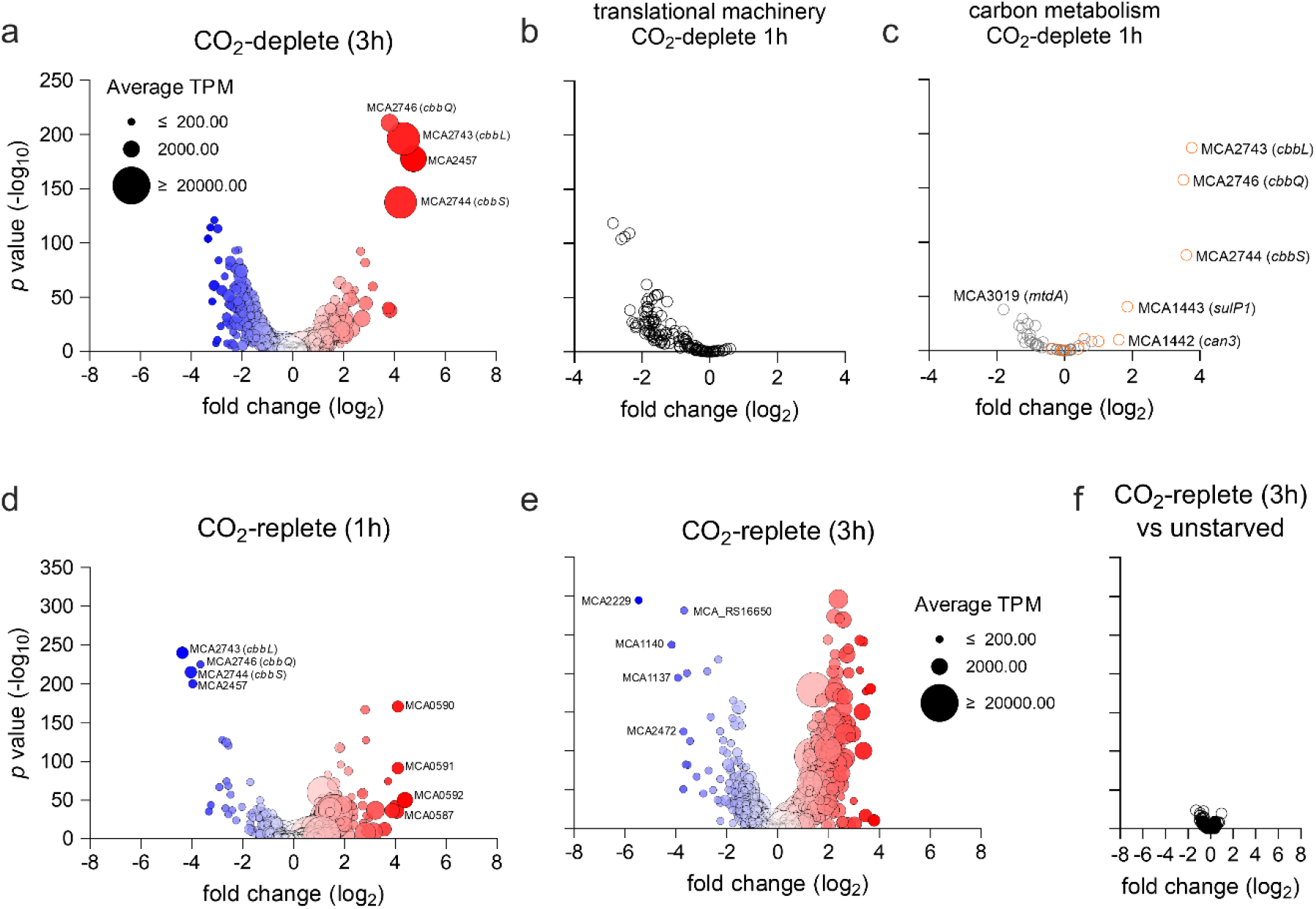
The *M. capsulatus* transcriptional response to CO_2_ starvation is reversed after CO_2_ supplementation. Volcano plots depicting (a) differentially expressed genes (DEGs) 3 h after CO_2_ removal from the bioreactor sparge gas stream (CO_2_-deplete) compared to unstarved (continuously sparged 20% CH_4_/2% CO_2_ in air) controls; a subset of DEGs encoding (b) translational machinery or (c) carbon metabolic enzymes 1 h after CO_2_-depletion; DEGs (d) 1 h and (e) 3 h after 2% CO_2_ was added to the sparge gas (CO_2_-replete) compared to CO_2_-deplete samples; and (f) DEGs in CO_2_-replete samples compared to unstarved controls. Genes in (b) were identified using GO-term annotations GO:0006412, GO:0006413, GO:0006414, and GO:0006415 and those in (c) were selected based on their involvement in CH_4_ (RuMP pathway, CH_4_ assimilation/dissimilation, oxidative pentose phosphate pathway) or CO_2_ (Calvin Benson Basham cycle, inorganic carbon concentrating) metabolism. In panels a, d, and e, red corresponds to upregulated genes and blue to downregulated genes with color intensity correlated to DEG log_2_ fold change. Dot size correlates to mean gene transcripts per million (TPM).

**Fig. S2.**
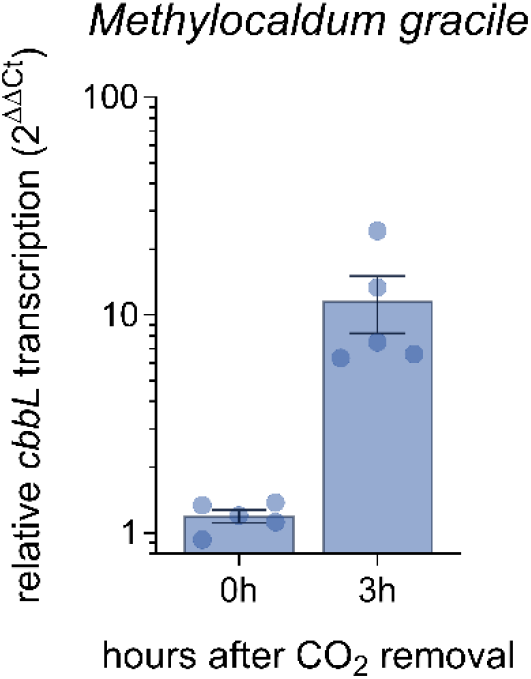
The Gammaprotobacteria Type Ib methanotroph, *Methylocaldum gracile* spp. *desertum* strain 0917 induces the *cbbL* gene encoding the RubisCO large subunit in response to CO_2_ starvation. RT-qPCR analysis of *cbbL* expression in cells cultivated with continuous gas sparging of 20% CH_4_/2% CO_2_ in air for 3 h followed by removal of the CO_2_ from the gas stream (20% CH_4_ in air, CO_2_-deplete). Methanotrophs were harvested for RNA extraction 3 h post CO_2_ starvation.

**Fig. S3.**
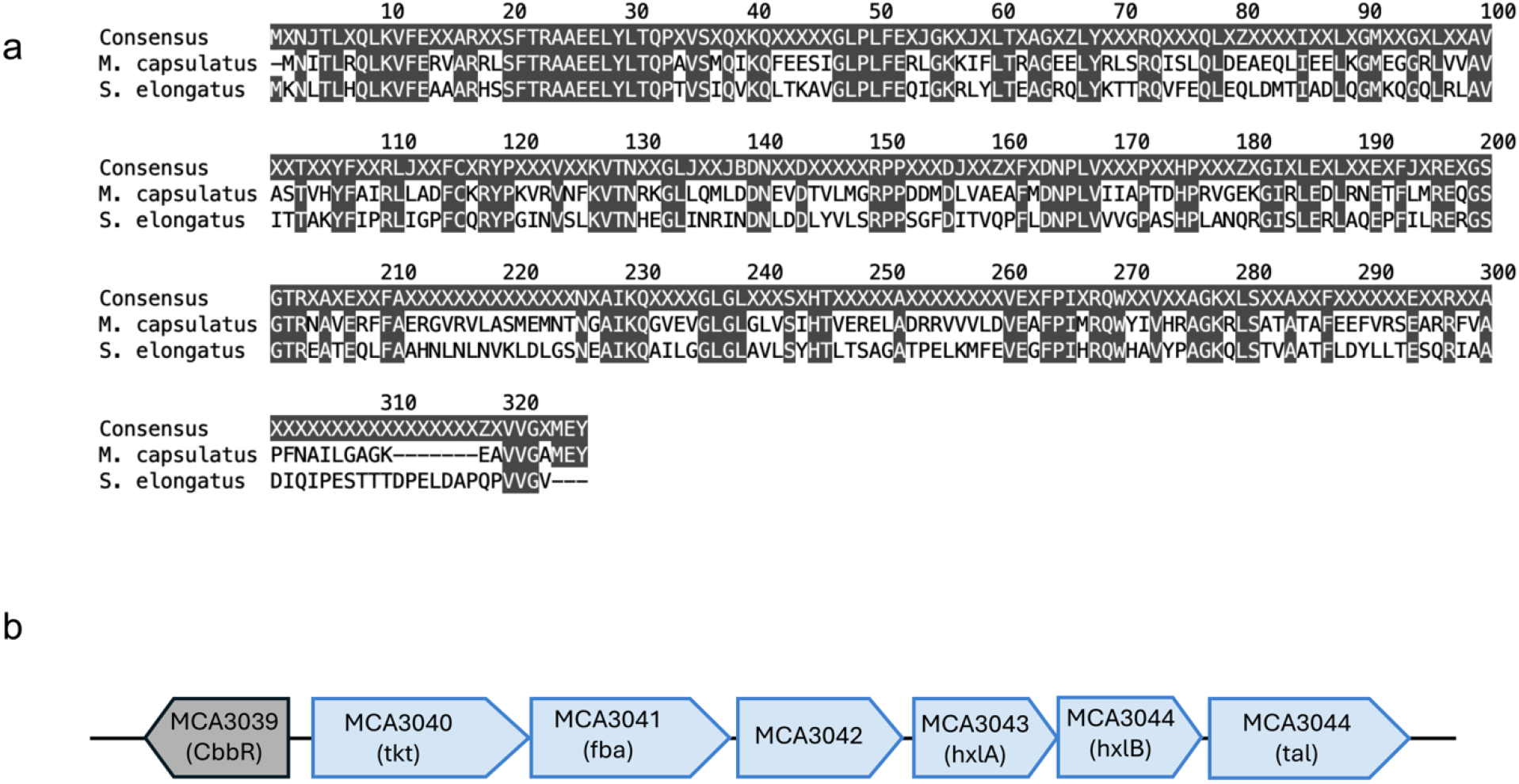
*Methylococcus capsulatus* CbbR alignment and genomic context. a) Clustal Omega alignment of the *M. capsulatus* CbbR homolog encoded by MCA3039 and CmpR from *Synechococcus elongatus* PCC 7942 (Synpcc7942_1310). Identical residues are highlighted. b) Schematic showing the genomic context of the CbbR-encoding gene MCA3039.

**Fig. S4.**
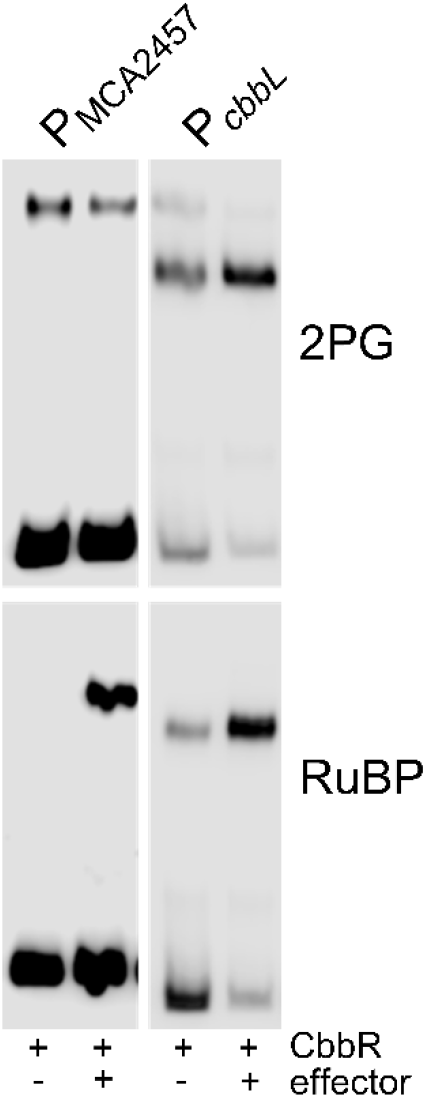
Supporting electromobility shift assays. a) CbbR-6xHis promoter binding EMSAs with 100 µM CbbR-6xHis and 1 mM effector molecule (2-phosphoglycolate (2PG), top; rubulose-1,5-bisphosphate (RuBP), bottom) and P_MCA2457_ (left) or P_cbbL_ (right) promoter probes.

**Other Supplementary Materials for this manuscript include the following:** Tables S1-S18 (separate excel file)

## Notes

### Competing Interest Statement

The authors have declared no competing interest.

